# The intracellular growth of the vacuolar pathogen *Legionella pneumophila* is dependent on the acyl chain composition of host membranes

**DOI:** 10.1101/2023.11.19.567753

**Authors:** Ashley A. Wilkins, Benjamin Schwarz, Ascencion Torres-Escobar, Reneau Castore, Layne Landry, Brian Latimer, Eric Bohrnsen, Catharine M. Bosio, Ana-Maria Dragoi, Stanimir S. Ivanov

## Abstract

*Legionella pneumophila* is an accidental human bacterial pathogen that infects and replicates within alveolar macrophages causing a severe atypical pneumonia known as Legionnaires’ disease. As a prototypical vacuolar pathogen *L. pneumophila* establishes a unique endoplasmic reticulum (ER)-derived organelle within which bacterial replication takes place. Bacteria-derived proteins are deposited in the host cytosol and in the lumen of the pathogen-occupied vacuole via a type IVb (T4bSS) and a type II (T2SS) secretion system respectively. These secretion system effector proteins manipulate multiple host functions to facilitate intracellular survival of the bacteria. Subversion of host membrane glycerophospholipids (GPLs) by the internalized bacteria via distinct mechanisms feature prominently in trafficking and biogenesis of the *Legionella*-containing vacuole (LCV). Conventional GPLs composed of a glycerol backbone linked to a polar headgroup and esterified with two fatty acids constitute the bulk of membrane lipids in eukaryotic cells. The acyl chain composition of GPLs dictates phase separation of the lipid bilayer and therefore determines the physiochemical properties of biological membranes - such as membrane disorder, fluidity and permeability. In mammalian cells, fatty acids esterified in membrane GPLs are sourced endogenously from *de novo* synthesis or via internalization from the exogenous pool of lipids present in serum and other interstitial fluids. Here, we exploited the preferential utilization of exogenous fatty acids for GPL synthesis by macrophages to reprogram the acyl chain composition of host membranes and investigated its impact on LCV homeostasis and *L. pneumophila* intracellular replication. Using saturated fatty acids as well as *cis*- and *trans*- isomers of monounsaturated fatty acids we discovered that under conditions promoting lipid packing and membrane rigidification *L. pneumophila* intracellular replication was significantly reduced. Palmitoleic acid – a C16:1 monounsaturated fatty acid – that promotes membrane disorder when enriched in GPLs significantly increased bacterial replication within human and murine macrophages but not in axenic growth assays. Lipidome analysis of infected macrophages showed that treatment with exogenous palmitoleic acid resulted in membrane acyl chain reprogramming in a manner that promotes membrane disorder and live-cell imaging revealed that the consequences of increasing membrane disorder impinge on several LCV homeostasis parameters. Collectively, we provide experimental evidence that *L. pneumophila* replication within its intracellular niche is a function of the lipid bilayer disorder and hydrophobic thickness.

## Introduction

Residence within host derived membrane-bound organelles is a broadly conserved survival strategy for many bacterial pathogens (Kumar and Valdivia, 2009; Manske and Hilbi, 2014; Schulz and Horn, 2015; Cornejo et al., 2017). Niche biogenesis and expansion are dynamic and involve remodeling of organelle-resident proteins as well as lipids through which trafficking to the correct cellular location and the maturation of the organelle into a compartment capable of nutrients acquisition is achieved (Toledo and Benach, 2015; Personnic et al., 2016; Weber and Faris, 2018; Allen and Martinez, 2020). These processes are generally driven by coordinated activities of multiple bacterial proteins translocated in the host cytosol by conserved bacteria-encoded secretion systems (Kubori and Nagai, 2016; Wagner et al., 2018; White and Cianciotto, 2019; Serapio-Palacios and Finlay, 2020). Host lipids regulate intracellular replication of vacuolar pathogens via multiple mechanisms: (i) as vacuole integrity guardians (Kumar and Valdivia, 2009); (ii) regulators of intracellular trafficking pathways (Cornejo et al., 2017); (iii) as a source of nutrients and building blocks for macromolecular biosynthesis (Carabeo et al., 2003; Toledo and Benach, 2015; Samanta et al., 2017; Huang et al., 2021); and (iv) as membrane tethering moieties for bacterial effector proteins (Weber et al., 2006; Ivanov et al., 2010; Hicks and Galán, 2013; Ivanov and Roy, 2013; Popa et al., 2016). Here, we explore how changes in the lipid composition of host membranes, in particular GPLs, impact the housing capacity of the vacuolar niche of the human intracellular pathogen *Legionella pneumophila* (Lp).

GPLs are membrane building-block lipids and key determinants of physiochemical properties of biological lipid bilayers (Meer et al., 2008). *De novo* GPL synthesis takes place at the ER membrane, where all the components are assembled by ER-localized enzymes (Jacquemyn et al., 2017). Conventional GPLs consist of a single polar head group and two hydrophobic acyl chains linked to the *sn*-1 and *sn*-2 positions of the glycerol backbone via ester bonds (Meer et al., 2008). The degree of saturation and carbon-chain length of the esterified fatty acids together with sterol content dictate lipid packing within membranes and thus impact membrane disorder, permeability, elasticity, curvature and fluidity (Lande et al., 1995; Bigay and Antonny, 2012; Ernst et al., 2016; Frallicciardi et al., 2022). In general, GPLs with saturated acyl chains pack tightly and rigidify membranes, whereas the opposite holds true for GPLs with unsaturated acyl chains. Eukaryotic organelles have distinct membrane GPL compositions and thus physical properties – for example, lipid disorder in organelle membranes increases inwards from the plasma membrane towards the ER (Bigay and Antonny, 2012; Jackson et al., 2016). Gradual decrease in the ratio of saturated (SFA) to monounsaturated acyl chains (MUFAs) within membrane GPLs is one-way the membrane disorder gradient from the plasma membrane (low) to the ER (high) is generated (Bigay and Antonny, 2012). Compared to other cellular organelles the ER membrane is in a liquid disordered phase and is relatively thin, both of these characteristics are key for its function as a proteostasis and lipogenesis hub (Erbay et al., 2009; Deguil et al., 2011; Halbleib et al., 2017; Jacquemyn et al., 2017).

In mammals, cells acquire lipids mainly through *de novo* synthesis as well as uptake from interstitial fluids or serum (Smith et al., 1978; Carvalho and Caramujo, 2018). Non-esterified fatty acids are present at various concentrations in human serum and represent a major lipid source for membrane biogenesis in cells (Abdelmagid et al., 2015; Alsabeeh et al., 2017; Jacquemyn et al., 2017). Internalized fatty acids are either (i) incorporated in newly synthesized GLPs for membrane biogenesis; (ii) stored in triglycerides; (iii) swapped with the acyl chains of preexisting membrane-resident GPLs or (iv) broken down via β-oxidation (McArthur et al., 1999; O’Donnell, 2022). The sequential action of phospholipases via the Land’s cycle allows for localized or global changes in membrane biophysical properties through the replacement of esterified saturated with unsaturated acyl chains or vice versa (O’Donnell, 2022). Influx of exogenous non-esterified fatty acids into cells results in rapid remodeling of membrane lipids dramatically affecting organelle function (Schaffer, 2003). One example is the activation of the ER stress response as a direct result of ER membrane rigidification caused by influx and incorporation of saturated fatty acids in ER-resident GPLs (Schaffer, 2003; Deguil et al., 2011). Acyl chain desaturation by the Stearoyl-CoA Desaturase (SCD) enzyme is one stress-resolving mechanism that restores lipid bilayer disorder and resolves ER membrane stress (Erbay et al., 2009; Piccolis et al., 2018; Oshima et al., 2020). Prolonged unresolved rigidification of the ER membrane results in rupture and ultimately cell death (Borradaile et al., 2006).

Lp is a pulmonary pathogen of the lower respiratory tract and the etiological agent of Legionnaires’ disease – a severe atypical pneumonia (Fraser et al., 1977; McDade et al., 1977). Lp replicates within a unique ER-derived membrane-bound compartment in mammalian macrophages referred to as the *Legionella*-containing vacuole (LCV) (Tilney et al., 2001). The maturation of the LCV into an organelle capable of supporting bacterial replication occurs within 4hrs post internalization (Ivanov and Roy, 2009) and is the result of coordinated action of over 300 Lp effector proteins translocated within the host cytosol by the T4bSS (Personnic et al., 2016; Cornejo et al., 2017). The *Legionella* T4bSS is a multi-component apparatus that is essential for pathogenesis and deletion of even a single gene encoding a structural element produces an avirulent mutant that fails to block endocytic maturation (Marra et al., 1992; Berger and Isberg, 1993; Andrews et al., 1998). A broadly conserved T2SS is also used by *Legionella* to secrete bacterial proteins within the lumen of the LCV (Truchan et al., 2017).

Lp manipulation of LCV-localized GPLs through a variety of mechanisms is central to trafficking, maturation and expansion of the bacteria-occupied organelle. Lp produced enzymes can modify GLPs headgroups (Hsu et al., 2012; Viner et al., 2012), one example of which is the generation of phosphoinositol-4-phosphate (PI4P) on the cytosolic leaflet of the LCV membrane that functions as an assembly platform for T4bSS effectors in part due to the prevalence of PI4P-binding domains with the Lp effector repertoire (Weber et al., 2006; Brombacher et al., 2009; Schoebel et al., 2010). The PI3P pool on nascent phagosomes occupied by Lp is converted into PI4P by the sequential activity of several host as well as Lp secreted phosphoinositide kinases and phosphatases, resulting in PI4P accumulation on the LCV that is atypical for the ER membrane (Swart and Hilbi, 2020).

In addition to posttranslational modifications, Lp can remodel membrane GPLs through the activity of different bacteria-encoded lipases secreted through the T2SS or T4bSS (Hiller et al., 2018). Phospholipases modify GPLs by hydrolyzing either (i) the carboxyl ester bond (PLA and PLB class) to produce lysophospholipids by removing an acyl chain from the *sn*-1 or *sn*-2 position; (ii) the glycerol-oriented phosphodiester bond (PLC class) to produce diacylglycerol; or (iii) the alcohol-oriented phosphodiester bond (PLD class) to produce phosphatidic acid by cleaving the headgroup (Aloulou et al., 2018). *Legionella* encodes at least 15 putative phospholipases with either PLA, PLC or PLD activities (Hiller et al., 2018), however with few exceptions their functionality in the context of infection is poorly understood. The T2SS effector PlaA has lysophospholipase as well as glycerophospholipid:cholesterol acyltransferase activities (Flieger et al., 2001, 2002; Banerji et al., 2005) and can destabilize the LCV membrane (Creasey and Isberg, 2012) if the host endosomal regulator inositol 5-phosphatase Oculocerebrorenal syndrome of Lowe 1 (OCRL1) is not removed from the compartment by the T4bSS effector SdhA (Choi et al., 2021). VipD - a patatin-like phospholipase T4bSS effector that localizes to early endosomes - removes PI3P through its PLA_1_ activity blocking endocytic maturation as a result (Gaspar and Machner, 2014; Lucas et al., 2014). Its paralog VpdC is a PLA_2_ lipase which when overproduced by Lp during infection increases cone-shaped lysophospholipids resulting in exacerbated membrane curvature and stunted LCV expansion (Li et al., 2022). Another family of Lp-encoded lipases (PlcA, PlcB & PlcC) that have been shown *in vitro* to have broad specificity phospholipase C activity are effectors of the T2SS (PlcA & PlcB) or the T4bSS (PlcC) that can hydrolyze GPLs to produce the signaling lipid 1,2-diacylglycerol (Aurass et al., 2013). A LCV localized T4bSS effector– LpdA – has been shown to remove the choline headgroup from phosphatidylcholine via its lipase D enzymatic activity generating phosphatidic acid in the process (Schroeder et al., 2015), which can be further dephosphorylated by the host phosphatidate phosphatase Lipin1 to generate diacylglycerols (Viner et al., 2012) providing evidence for GPLs headgroup remodeling taking place at the LCV membrane. Thus, extensive reprogramming of host GPLs by *Legionella*-encoded enzymes is one multifaceted mechanism through which niche biogenesis and homeostasis is achieved.

LCV expansion is also controlled in part by *de novo* lipogenesis (Abshire et al., 2016; Ivanov, 2017), driven by the host metabolic checkpoint serine/threonine kinase Mechanistic Target of Rapamycin (MTOR). MTOR integrates sensory cues from energy and nutrient pathways to ensure fidelity and coordination of anabolic processes in eukaryotic cells (Abshire et al., 2016; Shimobayashi and Hall, 2016). Lp activates MTOR signaling downstream of the amino-acid sensing pathway (Abshire et al., 2016; Leon et al., 2017) in infected macrophages by increasing the amount of intracellular free amino acids through a blockade in host protein translation elicited by T4bSS effectors (Leon et al., 2017). Consequently, MTOR inhibition limits membrane expansion causing LCV rupture, which can be overcome by supplementation with exogenous cholesterol and other serum-derived lipids (Abshire et al., 2016).

Considering the morphological changes as well as the extensive molecular remodeling of the LCV membrane during its lifespan, it is important to determine to what extend niche homeostasis is regulated by the acyl chain composition of the membrane GPLs, especially because of their crucial structural role in biological membranes. Here, we investigate the role of esterified fatty acids in Lp intracellular replication in macrophages.

## Results

### Dietary fatty acids alter the capacity of Legionella-infected macrophages to support bacterial replication but do not function as a growth-promoting nutrient for Legionella pneumophila

We were interested in investigating to what extend variations in membrane lipid composition impact Lp infection and reasoned that one way to approach this question experimentally is though metabolic reprogramming of the GPL repertoire via exogenous lipid supplementation because eukaryotic cells preferentially utilize fatty acids internalized from interstitial fluids (exogenous dietary fatty acids) over *de novo* synthesized lipids for GPL assembly. To this end, we infected primary bone-marrow derived murine macrophages (BMMs) with Lp and measured bacterial growth over several infection cycles under conditions where different fatty acids from the two major membrane classes (saturated and monounsaturated) (Table 1) were supplemented in the culture media for the duration of the infection (Fig 1A). These infections were carried out under serum-free conditions to ensure preferential utilization by macrophages of the specific fatty acid tested over other serum-derived lipids. For these assays, the fatty acids were complexed with albumin, which is the main transport carrier of non-esterified fatty acids in interstitial fluids (Spector, 1975; VUSSE, 2009). A bioluminescent Lp strain encoding the *luxR* operon under a constitutive promoter was used which allowed us to monitor bacterial growth continuously in high throughput manner (Ondari et al., 2022).

**Figure 1.**
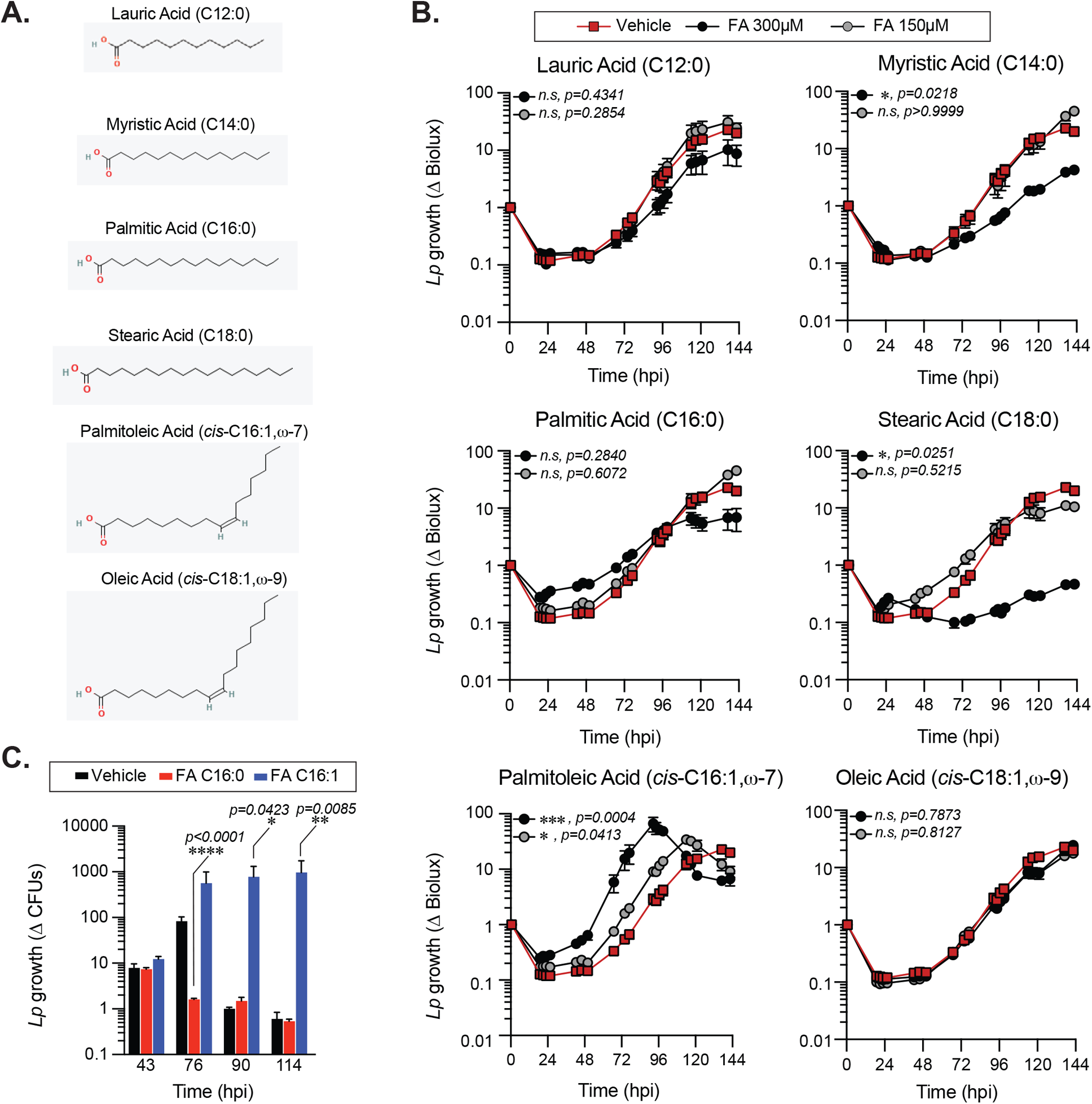
Exogenous Fatty acids alter Lp growth kinetics in primary BMMs infections. **(A)** 2-D chemical structures indicating the size and saturation state of the fatty acids used in the experiments. Oxygen and hydrogen atoms are highlighted in red and green, respectively. Images were sourced from PubChem (pubchem.ncbi.nlm.nih.gov) **(B-C)** Lp growth kinetics with primary BMMs at MOI=2.5 as measured by fold-change in bioluminescence signal **(B)** or recovery of colony forming units **(C)** from T_0_. The culture media was supplemented with different fatty acids bound to BSA at the indicated concentrations or with BSA alone (vehicle control). Average ± StDev of three technical replicates are shown. Each data series from treated-cells was compared to vehicle-treated control using ordinary one-way ANOVA analysis and the respective p-values are indicated. At least three biological replicates for each experiment were generated. (*n.s* – not significant, FA C16:0 – palmitic acid, FA 16:1 – palmitoleic acid).

**Table 1.** Fatty acids used in this study. Saturated and monounsaturated fatty acids ranging from 12 to 18 carbons in length were used. The double bonds position from the methyl group of the fatty acid (⍵) and the geometrical orientation (*cis* vs. *trans*) are shown.

| Common Name | Nomenclature | Double bonds |  |
| --- | --- | --- | --- |
|  |  | position | orientation |
| Lauric Acid | C12:0 | - | - |
| Myristic Acid | C14:0 | - | - |
| Palmitic Acid | C16:0 | - | - |
| Palmitoleic Acid | *cis*-C16:1, ω-7 | ω-7 | *cis* |
| Sapienic Acid | *cis*-C16:1, ω-10 | ω-10 | *cis* |
| Palmitelaidic Acid | *trans*-C16:1, ω-7 | ω-7 | *trans* |
| Stearic Acid | C18:0 | - | - |
| Oleic Acid | *cis*-C18:1, ω-9 | ω-9 | *cis* |
| Elaidic Acid | *trans*-C18:1, ω-9 | ω-9 | *trans* |

When supplied exogenously myristic (C14:0), stearic (C18:0) and palmitoleic acid (C16:1) significantly altered Lp intracellular growth in primary BMMs in a dose-dependent manner (Fig 1B). Overall, all SFA tested reduced Lp growth in primary BMMs infections, although only the differences for myristic and stearic acid treatments were statistically significant (Fig 1B). Conversely, MUFAs either significantly enhanced Lp growth during infection (palmitoleic acid) or did not produce a growth phenotype (oleic acid) (Fig 1B). An alternative assay measuring the colony forming units (CFUs) recovered at distinct times after infection also produced similar bacterial growth enhancement by palmitoleic acid (Fig 1C). Comparable Lp growth patterns were seen in infections of immortalized BMMs (iBMMs) where palmitoleic acid enhanced and all of the SFA except lauric acids (C12:0) reduced bacterial growth (SFig 1A-B). The mean amount of circulating palmitoleic acid in the serum in a large cohort of 827 young healthy adults from diverse ethnic backgrounds was measured at 133.0 ± 67.2 µM (min 27.7µM and max 555.9µM) (Abdelmagid et al., 2015). Therefore, we conclude that palmitoleic acid supplied at physiologically relevant concentrations can promote Lp growth in infections of primary and immortalized BMMs.

The growth-altering phenotypes produced by dietary fatty acids in Lp infections could be a result of a nutrient effect due to direct metabolic utilization by the bacterium, which should be evident when bacterial growth is measured in the absence of host cells. Therefore, we measured Lp growth in AYE liquid cultures in the presence of different fatty acids or vehicle control condition. The initiation of Lp replication was delayed in axenic cultures in the presence of 300µM lauric, myristic, palmitic or oleic acids to various degrees (Fig 2A), indicating that these fatty acids can induce measurable direct growth defect in the absence of host cells. Under the same growth conditions *Escherichia coli* growth in axenic media was enhanced (Fig 2B) consistent with *E. coli* capacity for metabolic utilization of exogenous fatty acids (Jaswal et al., 2021). Palmitoleic acid at concentrations that enhanced Lp intracellular replication did not affect Lp growth in axenic cultures (Fig 2A). Thus, palmitoleic acid in the absence of host cells is not a growth-promoting metabolite for *Legionella*. We also conclude that Lp tolerance to fatty acids-induced lipotoxicity is reduced compared to *E.coli*.

**Figure 2.**
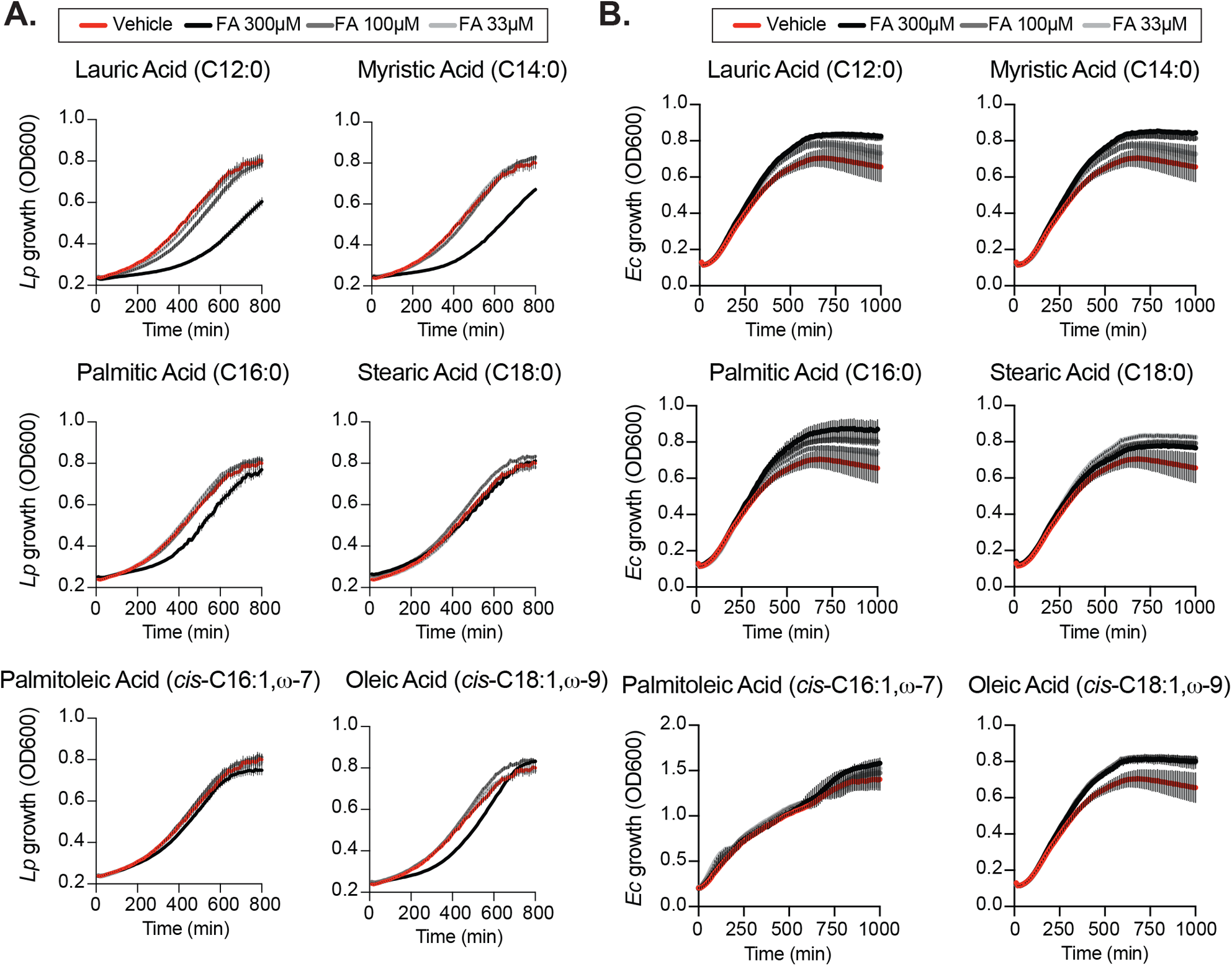
Effects of exogenous fatty acids on bacterial growth in axenic cultures. **(A-B)** Axenic growth kinetics of Lp **(A)** and *E. coli* **(B)** as measured by changes in optical density (OD_600_) over time in liquid cultures supplemented with increasing concentrations (33 to 300µM) of the indicated fatty acids bound by BSA or with BSA alone (vehicle control). OD_600_ data was collected every 10 minutes and each data series represents an average of three technical replicates ± StDev. One representative of three biological replicates is shown for each experiment.

### Dietary palmitoleic acid increases macrophage permissiveness for Lp replication by modulating multiple LCV homeostasis parameters

For *Legionella*, the number of bacteria egressing from each infected macrophage is determined by several variables – (i) internalization kinetics; (ii) trafficking and remodeling of the LCV into an ER-like compartment; (iii) maturation of the LCV into an organelle that supports bacterial replication; (iv) synchronization of LCV expansion and bacterial replication and (v) LCV membrane disruption during egress. Even moderate variations in these key parameters can be amplified over successive infection cycles to produce a growth phenotype. We sought to determine which variable(s) is/are directly or indirectly regulated by the presence of exogenous palmitoleic acid.

First, we compared the uptake of BSA-coupled fatty acids by BMMs as different rates of internalization could potentially explain the distinct Lp growth phenotypes we observed during infection. Fatty acids uptake occurs through direct membrane diffusion of the lipid or facilitated transport by membrane transporters (Su and Abumrad, 2009), where the transport rates depend on size and saturation states of the fatty acid (Kampf et al., 2006; LeBarron and London, 2016). One fate of internalized fatty acids is the rapid re-esterification into newly synthesized triacylglycerols for storage into lipid droplets (Walther and Jr., 2012). Thus, we quantified the number of lipid droplets per cell as an indirect readout for fatty acids uptake by iBMMs (Fig 3). After a 4hrs treatment all fatty acids increased the lipid droplets’ size (Fig 3A) and significantly increased the number of lipid droplets per cell (Fig 3A-B) consistent with increased internalization and incorporation of dietary fatty acids in triacylglycerols. Approximately, a two-fold increase in the number of lipid droplets was observed in MUFAs-treated iBMMs compared to SFA-treated cells potentially as a result of higher transport rates (Fig 3B). Nevertheless, an increased lipid uptake is unlikely to account for the Lp growth phenotype caused by dietary palmitoleic acid because all three MUFAs induced lipid droplet formation comparably (Fig 3B), yet Lp growth enhancement was specific for palmitoleic acid (Fig 1B and SFig 1A).

**Figure 3.**
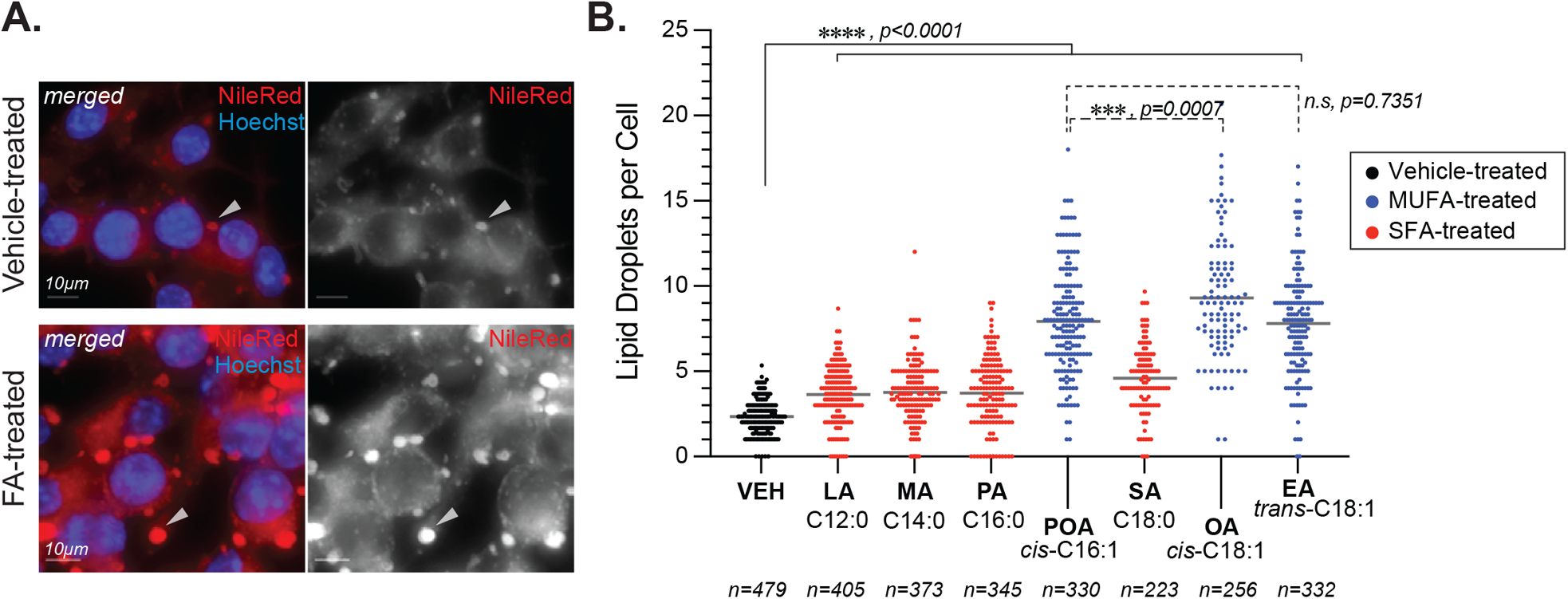
Lipid droplets formation in fatty acids-treated macrophages. **(A-B)** Fatty acids supplementation increases the number of lipid droplets in the cytosol of treated iBMMs. **(A)** Representative micrographs showing neutral lipids accumulation in lipid droplets (arrowheads) in the cytosol of iBMMs treated with a fatty acid for 4hrs at 150µM or with vehicle alone. Cells were stained with the neutral lipids dye NileRed that accumulates in lipid droplets. Scale bar = 10µm. **(B)** Quantitation of the number of lipid droplets per cell in iBMMs treated with the indicated saturated (SFA) or monounsaturated fatty acids (MUFAs). The number of cells analyzed in each condition is indicated. Ordinary one-way ANOVA was used for statistical comparison among the data series and p-values for each comparison are indicated. Representative data from one of two biological replicates are shown. LA - *lauric acid*; MA - *myristic acid*; PA - *palmitic acid*; SA - *stearic acid*; POA - *palmitoleic acid*; EA - *elaidic acid*

Next, we investigated whether Lp phagocytic uptake was enhanced by dietary palmitoleic acid, which may account for the increased bacterial growth seen in the BMM infection assays. To this end, we pre-treated iBMMs with palmitoleic acid or vehicle control for 16hrs after which *Lp* internalization was measured at 2hpi using inside/out microscopy approach (Fig 4A-B). Palmitoleic acid treatment did not alter the number of internalized bacteria by iBMMs, unlike the disruption of actin polymerization by cytochalasin D which interfered with phagocytosis (Fig 4B). Therefore, we conclude that an increase in bacteria uptake is not the cause for the higher bacterial growth supported by palmitoleic acid-laden BMMs.

**Figure 4.**
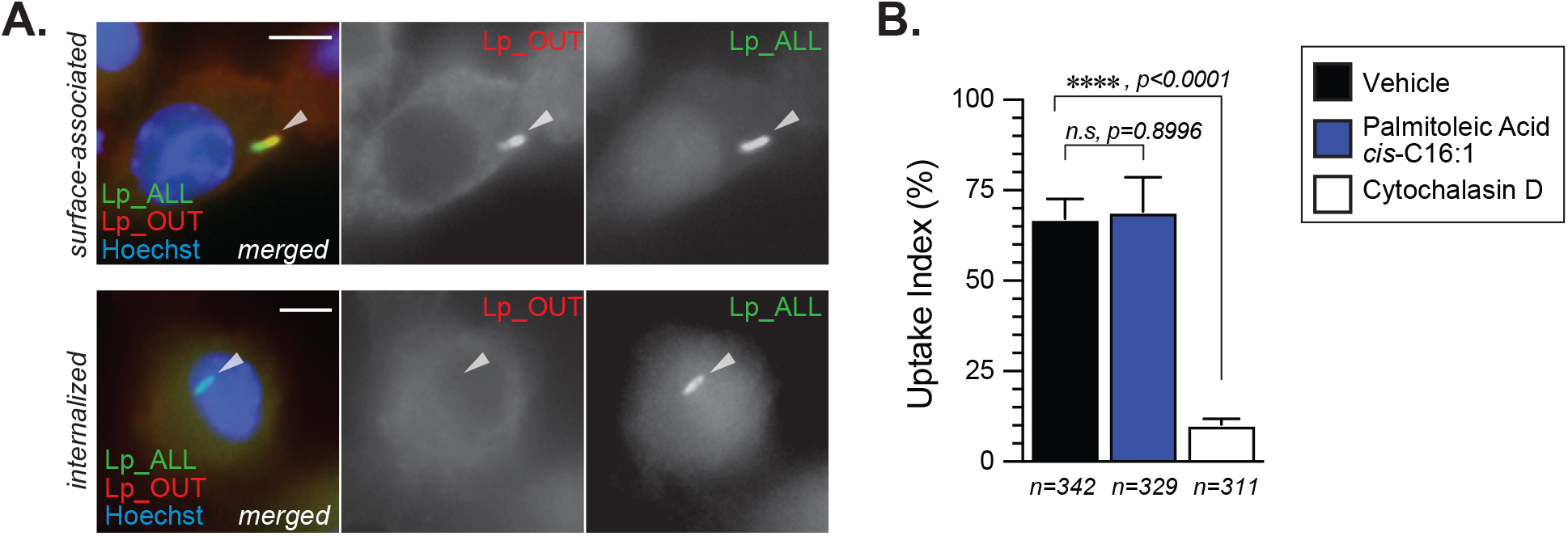
Lp internalization by iBMMs is not altered by palmitoleic acid supplementation. **(A-B)** Data shows bacteria uptake by macrophages as measured by microscopy with the inside/out staining approach. **(A)** Representative micrographs of iBMMs infected by GFP-expressing Lp that were stained with anti-Lp antibody without permeabilization showing single-positive internalized (bottom panels) and dual-positive surface-associated bacteria. Scale bar = 5µm. **(B)** Quantitative analysis of Lp internalization at 2hpi by iBMMs that were pretreated for 16hrs with either 300µM palmitoleic acid bound by BSA or with vehicle control only. Control condition in which phagocytosis is blocked by the pre-treatment with the actin polymerization inhibitor cytochalasin D (5µM) is also shown. Data represents the average of three biological replicates ± StDev where >100 bacteria for each treatment were scored. Total number of cells analyzed for each condition is shown. For statistical analysis, one-way ANOVA was used and the p-value for each comparison is shown.

Because *Lp* uptake was not enhanced by dietary palmitoleic acid, we next investigated the effect on LCV biogenesis and expansion. Lp replication is initiated in mature LCVs at ∼ 4 hours post internalization, after trafficking and remodeling of the LCV into an ER-like organelle is completed (Ivanov and Roy, 2009). While majority of Lp phagocytosed by BMMs successfully evade endosome maturation (> 90%) (Roy et al., 1998) and the organelle remodeling is successfully completed, only ∼50% of mature LCVs support robust bacterial replication (Nogueira et al., 2009). Therefore, we investigated whether dietary palmitoleic acid increases the number of LCVs supporting bacterial replication which could account for the growth-enhancement phenotype. To this end, we used a well-established microscopy-based assay to calculate the percentage of LCVs supporting bacterial replication (Nogueira et al., 2009). In this method the percentage of LCV supporting bacterial replication is derived as an index of the percentage of infected cells harboring LCVs that support bacterial replication at 10hpi over the percentage of infected cells at 2hpi (Fig 5). In general, the LCVs in palmitoleic acid-laden iBMMs at 10hpi harbored more bacteria (Fig 5A-B) with fewer vacuoles containing less than 3 bacteria as compared to vehicle control (7% vs 23% in vehicle control) (Fig 5B). The RV index also reflected the capacity of palmitoleic acid to increase significantly the percentage of LCVs supporting bacterial replication from 58.3% to 81.5% (Fig 5C). Moreover, the percentage of large LCVs (bacteria n >13) in palmitoleic acid treated iBMMs doubled from 35% to 71% indicating an overall increased LCV capacity to support bacterial replication. Based on these data we conclude that dietary palmitoleic acid (i) enhances the capacity of LCVs to support Lp replication and (ii) increases the number of LCVs that support bacterial replication.

**Figure 5.**
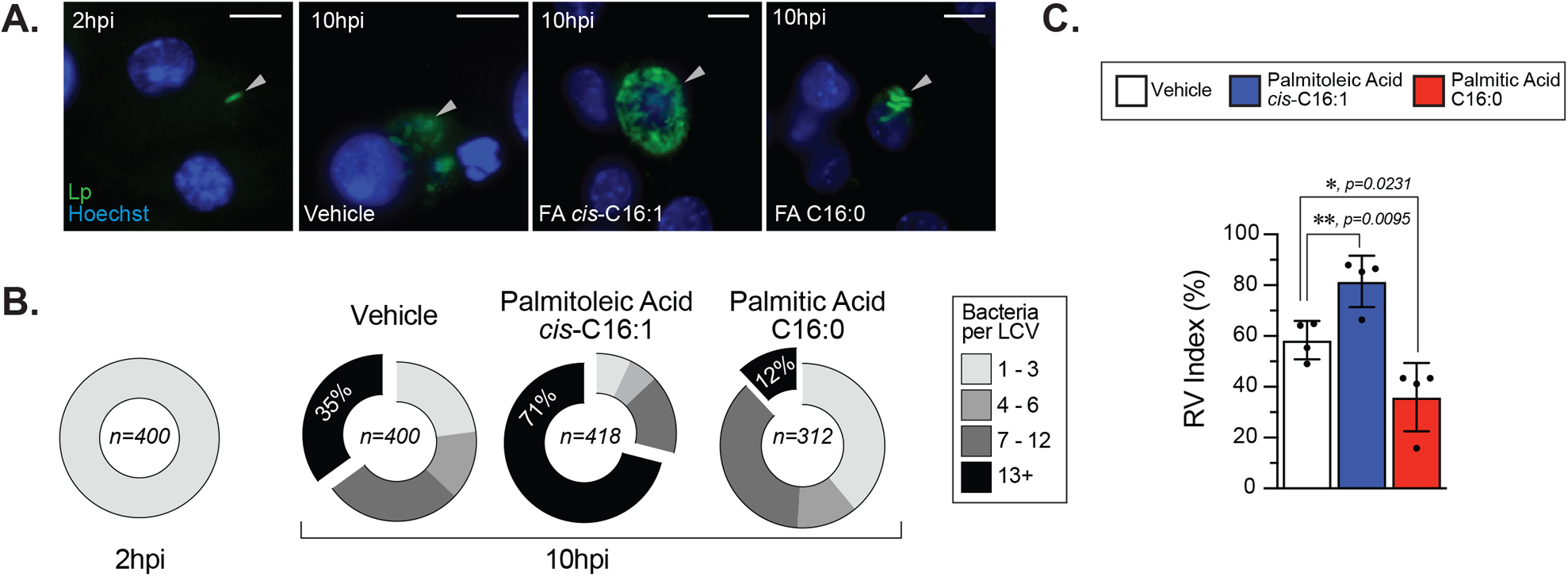
Palmitoleic but not palmitic acid increases the percentage of LCV that support bacterial replication. **(A-C)** Microscopy analysis of Lp intracellular replication at 10hpi in iBMM infections where cells were treated with either the indicated BSA-bound fatty acids (300µM) or BSA only control starting at 2hpi. **(A)** Micrographs showing representative Lp-containing vacuoles harbored by iBMMs infected with GFP-expressing Lp for 10 hours. Scale bar = 10µm. **(B)** Graphed data shows binned size distribution of LCVs harbored by iBMMs at 10hpi after the cells were treated as indicated. Data is pooled from four biological replicates and the number of LCVs analyzed for each condition is indicated. **(C)** The Replicate Vacuole (RV) Index indicates the percentage of LCVs that support bacterial replication at 10hpi and was calculated by dividing the percentage of infected cells harboring LCVs that support bacterial replication (i.e. contain >6 bacteria) by the percentage of infected cells at 2hpi and the result was multiplied by 100. Averages of four biological replicates ± StDev are shown where each data point is derived from a distinct biological replicate. At least 78 LCVs from each condition per biological replicate were analyzed and used for the RV Index calculation. For calculation of statistical significance, each data series was compared to the data from vehicle-treated cells using a one-way ANOVA paired analysis and the p-values are indicated.

To gain insight into which aspects of LCV homeostasis are impacted by different fatty acids we performed live-cell microscopy of BMMs infected with GFP-expressing Lp, where cells were imaged every 4 hours over a 2-day period (Fig 6). Using this approach, three distinctly identifiable phases were observed during the LCV lifespan – expansion, stasis and collapse. During the *expansion* phase the size of the LCV increases, whereas during the *stasis* phase little measurable change in the LCV size was detected (Fig 6A). The *collapse* phase of the LCV is initiated by an abrupt drop in LCV size accompanied by sudden decrease in intensity of the GFP signal consistent with membrane rupture and is terminated when the GFP signal disappears entirely indicating egress completion (Fig 6A). Bulk analysis at the population level revealed that in palmitoleic acid-treated cells the LCV size on average increased significantly faster as compared to vehicle or palmitic acid-treated cells even though comparable number of LCVs were detected across all conditions for the first 20 hours of the infection cycle (Fig 6B), which suggest that LCV expansion might be affected. However, from averaged population data of an asynchronous infection we could not determine whether this phenotype is due to earlier initiation of bacterial replication, faster bacterial replication or prolonged LCV lifespan. Therefore, we tracked the size of individual LCVs over their lifespan from each experimental condition to determine the maximum LCV size (Fig 6D), the duration of the LCV expansion phase (Fig 6E), the entire LCV lifespan (Fig 6F) and the LCV rate of expansion (Fig 6G). From single particle tracking analysis, we observed that in palmitoleic acid-laden macrophages the duration of LCV expansion on average was prolonged (20hrs vs 16hrs in vehicle control) (Fig 6E), the LCV reached a larger peak size (295.4 µm^2^ vs 205µm^2^ in vehicle control) (Fig 6D), and the LCVs doubled in size quicker (in 5.1 hours vs 6.1 hours for vehicle control) (Fig 6G). The individual size traces of 10 randomly selected LCVs showed that bacterial replication did not initiate earlier in palmitoleic acid-laden macrophages compared to vehicle control treated cells (SFig 2). In addition, palmitoleic acid treatment shortened the LCV collapse phase resulting in higher percentage of LCVs that broke down within 4hrs (19% vs 6% in vehicle control) (SFig 3). Thus, we conclude that palmitoleic acid treatment remodels macrophages in a manner that makes them more permissive for Lp growth by prolonging the LCV expansion phase and accelerating bacterial replication rather than triggering early bacterial replication. In addition, we found evidence for shortened LCV collapse phase, which likely accelerated egress and the initiation of the subsequent infection cycle. The LCV membrane is one element common to all these phenotypes, suggesting that remodeling of host membrane lipidome by palmitoleic acid may be at the center.

**Figure 6.**
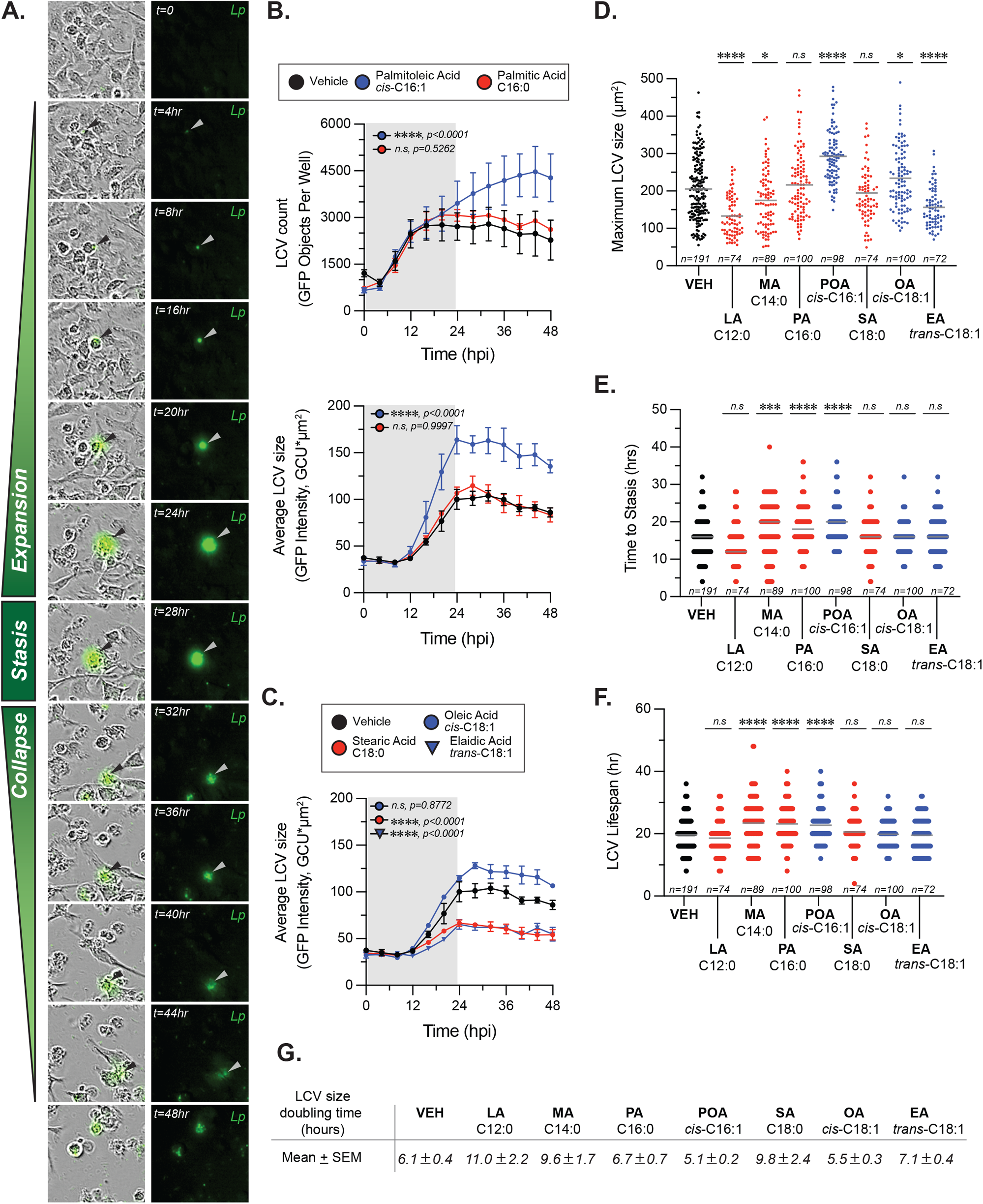
Exogenous fatty acids alter LCV homeostasis in infections of primary BMMs. **(A-F)** Live-cell imaging of Lp intracellular replication in primary BMMs infected with GFP-expressing bacteria at MOI=5. BSA-complexed exogenous fatty acids (150µM) were added at 2hpi. **(A)** The sequence of micrographs tracks the expansion, stasis and collapse of a representative LCV over a 44-hour period. Merged (left panels) and fluorescence (right panels) micrographs are shown. **(B-C)** Population level analysis of LCV dynamics during infection in the presence of exogenous BSA-complexed fatty acids or BSA only vehicle control. In **(B)** the top panel graph tracks the average number of LCVs over time, whereas the bottom panel graph in **(B)** and the data in **(C)** tract the average LCV size over time. Averages ± StDev of three technical replicates are shown. **(D-F)** Quantitative analysis of LCV dynamics at the level of individual LCVs. For each condition, the growth parameters of at least 72 individually tracked LCVs were analyzed. Each data point in the graph is derived from a single vacuole. The maximum size reached by individual LCVs under the indicated infection conditions are shown in **(D)**. The length of the expansion phase for each vacuole is plotted in **(E)** and the total LCV lifespan is graphed in **(F)**. **(G)** The area doubling time ***G*** for each LCV was calculated as follows ***G*** =(T_MAX_-T_0_)/(3.3*log (LCV_SIZE_ [max])/(LCV_SIZE_ [T_0_])), where T_MAX_ indicates the time LCV reaches maximum size and T_0_ is the time the LCV is first observed. The mean LCV doubling time ± SEM is shown for each treatment. **(B-F)** Each data series from treated cells was compared to the vehicle-treated control using ordinary one-way ANOVA analysis and the respective p-values are indicated. Representative data from one of three biological replicates are shown. (*n.s* – not significant). LA - *lauric acid*; MA - *myristic acid*; PA - *palmitic acid*; SA - *stearic acid*; POA - *palmitoleic acid*; EA - *elaidic acid*

### Dietary palmitoleic acid is incorporated in specific glycerophospholipids during infection

To gain insight into the remodeling of the macrophage lipidome during *Legionella* infection mediated by dietary palmitoleic acid we performed lipidomics analysis. To this end, primary BMMs were uninfected or infected for 8hrs in the presence/absence of 200µM palmitoleic acid followed by bulk lipids extraction and liquid chromatography tandem mass spectrometry analysis (Fig 7). The lipid abundance for 555 distinct lipids across all experimental conditions was determined. Both C16:0 and C16:1 acylated glycerophospholipids increased upon Lp infection, however palmitoleic acid supplementation further increased significantly the amount of C16:1 acylated GPLs but not C16:0 acylated GPLs demonstrating the internalization and specific incorporated of dietary palmitoleic acid into membrane glycerophospholipids (Fig 7A). Preferential incorporation of C16:1 acyl chains was detected in phosphatidylcholines (PC) as well as phosphatidylglycerols (PG) and to a lesser extend in phosphoethanolamines (PE) and phosphatidylserines (PS) (Fig 7A). GPLs containing C16:1 acyl chain at either *sn-1* or *sn-2* positions were enriched following palmitoleic acid supplementation indicating certain membrane GPLs are also remodeled via Land’s cycle through direct acyl chain substitution involving PLA_1_ and PLA_2_ phospholipases (O’Donnell, 2022) (Fig 7A). Acyl chain analysis on the top 100 lipids significantly changed in response to infection revealed that palmitoleic acid treatment caused also notable decrease in C18:0 acylated GPLs specifically (Fig 7B), which indicates potential replacement of C18:0 acyl chains by exogenous C16:1. The consequence of exchanging saturated C18:0 with monounsaturated C16:1 on membrane GPLs is an overall increase in the degree of unsaturation resulting in enhanced membrane disorder, fluidity and permeability. Increased membrane fluidity at the ER increases ER tolerance to proteostatic stress and membrane rigidification-induced lipotoxicity (Erbay et al., 2009; Jacquemyn et al., 2017).

**Figure 7.**
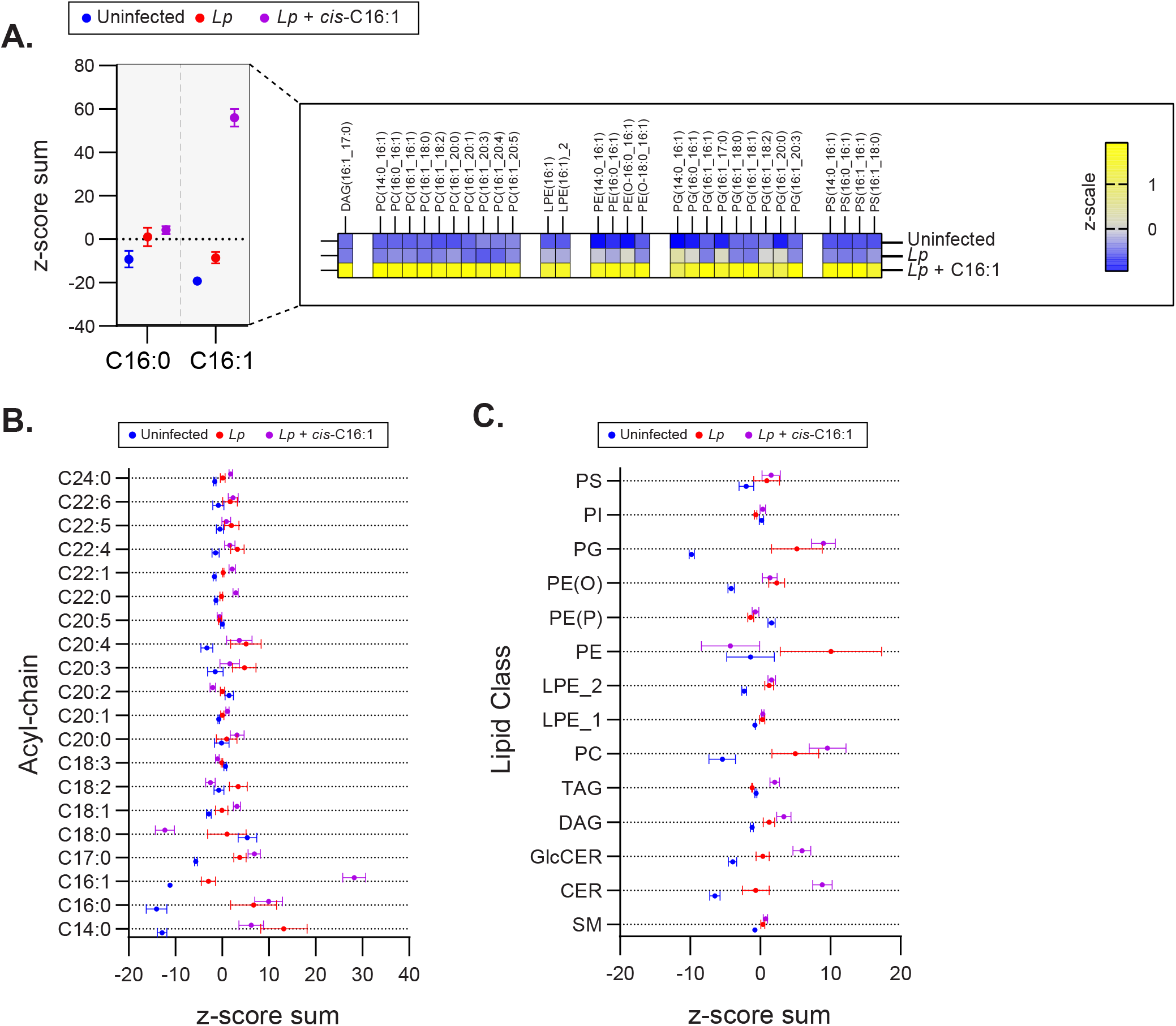
Exogenous palmitoleic acid is incorporated in membrane glycerophospholipids during macrophage infections with Lp. **(A-C)** Lipidomics comparison between uninfected primary BMMs and BMMs infected by Lp for 8hrs in the presence or absence of 200µM palmitoleic acid (C16:1) complexed with BSA. **(A)** Data in the left panel shows the z-score sum of the C16:1-containing lipids among the top 100 lipids that changed significantly in uninfected vs Lp + C16:1 condition as determined by Student’s t-test. The z-score sum of palmitoleic acid (C16:0)-containing glycerophospholipids is included as control to illustrate the highly specific accumulation of C16:1-containing lipids. All C16:1-containing lipids that make up the z-score sum are broken down in the right panel in a heatmap of the average z-scale by condition and show that supplementation of exogenous C16:1 results in direct integration into mainly PC and PG lipids and to a lesser extent in PE and PS lipids. **(B-C)** Breakdown of the top 100 lipids that changed significantly in uninfected vs Lp infection conditions by acyl-chain composition **(B)** and lipid class **(C)**. **(A-C)** The analyzed data was obtained from eight biological replicates. PC – *phosphatidylcholines*; PG – *phosphatidylglycerols*; PE – *phosphoethanolamines*; PS-*phosphatidylserines*; PI – *phosphoinositides*; PE (O) - *plasmanyl ether-linked PEs*; PE (P) - *plasmenyl ether-linked PEs*; LPE_2 – *sn-2 acylated lyso-phosphatidylethanolamines*; LPE_1 – *sn-1 acylated lyso-phosphatidylethanolamines*; TAG – *triacylglycerols*; DAG – *diacylglycerols*; GlcCER – *glucosylceramides*; CER – *ceramides*; SM – *sphingomyelins*

Dietary palmitoleic acid also altered the infection-induced enrichment of certain membrane lipid classes (Fig 7C). For example, PC and ceramides were further enriched by palmitoleic acid treatment, whereas the enrichment of PE was reversed (Fig 7C). Predictably the levels of di- and triacylglycerols, which can store fatty acids, also increased likely to accommodate the influx of exogenous palmitoleic acid (Fig 7C). Taken together, the data indicates that exogenous palmitoleic acid in Lp-infected macrophages remodels multiple aspects of the host lipidome one of which – replacement of C18:0 with C16:1 in membrane GPLs through preferential incorporation in PC and PG – would increase membrane disorder and fluidity. Potentially, membrane disorder could be an important parameter for sustained optimal LCV membrane expansion.

### Lp intracellular replication in the presence of exogenous fatty acids is a function of the carbon chain size, saturation state and double-bond position in monounsaturated fatty acids

The lipidomics data prompted us to investigate whether alteration in membrane disorder can account for the growth-enhancing capacity of palmitoleic acid. The Lp intracellular growth defects induced by supplementation of saturated fatty acids, which rigidify membranes upon GPLs incorporation, also are consistent with that idea (Fig 1B and SFig 1A). In general, the negative impact of exogenous SFAs on LCV homeostasis is evident from the smaller LCV size (Fig 6 C-D), the decrease in the rate of LCV expansion (Fig 6G) and the prolonged egress (SFig 3). However, at higher concentrations SFA exhibited some lipotoxicity to both Lp (Fig 2A) and BMMs (SFig 4A); thus, SFA treatment produces multiple phenotypes in both macrophages and Lp making it difficult to attribute the growth restriction phenotype solely to LCV expansion. Hence, we took a different approach by taking advantage of *cis*-*trans* geometric and positional isomerism of MUFAs to explore the role of membrane disorder in Lp intracellular growth. The *trans-*MUFA isomers have linear structures similar to SFAs (Figs 1A and 8A) and like SFAs increase membrane rigidification when incorporated in GPLs (Deguil et al., 2011; Tyler et al., 2019). Palmitelaidic acid (*trans*-C16:1, μ-7) – the geometric isomer of palmitoleic acid (*cis*-C16:1, μ-7) - reduced Lp intracellular growth in iBMM infections in a dose-dependent manner when supplied exogenously (Fig 8B). Importantly, palmitelaidic acid did not impact Lp growth under axenic conditions (Fig 8C) demonstrating that this lipid lacked microbiocidal activity and the growth phenotype is likely a result of lipidome reprogramming of the host cell. Similarly, elaidic acid (*trans*-C18:1, μ-9) - the linear geometric isomer of oleic acid (*cis*-C18:1, μ-9) – also produced a significant Lp growth defect phenotype in macrophage infections in a dose-dependent manner whereas oleic acid treatment had a neutral effect on Lp growth in biolux assays (Fig 8B) and a small but measurable increase in LCV size in live-cell imaging assays (Fig 6D). Macrophages exposed to elaidic acid harbored smaller LCVs as compared to oleic acid-treated macrophages (Fig 6C-D), which was a result of decreased LCV rate of expansion (Fig 6G) and did not affect the LCV lifespan (Fig 6F). Elaidic acid did not reduce macrophage viability (SFig 4B) and had only a minor impact on Lp growth in axenic cultures at high concentrations (Fig 8C). Taken together the data from treatments with matched positional isomers demonstrates that membrane rigidifying *trans*-MUFAs interfere with LCV expansion and Lp growth unlike their geometric non-linear *cis*-isomers which increase membrane disorder. Thus, we conclude that the physiochemical properties especially pertaining to membrane disorder dictate the optimal rate of niche expansion and housing capacity of the LCV. The distinct Lp growth phenotypes elicited by palmitoleic vs oleic acid indicate that the carbon chain length of the fatty acid is also a determinant.

**Figure 8.**
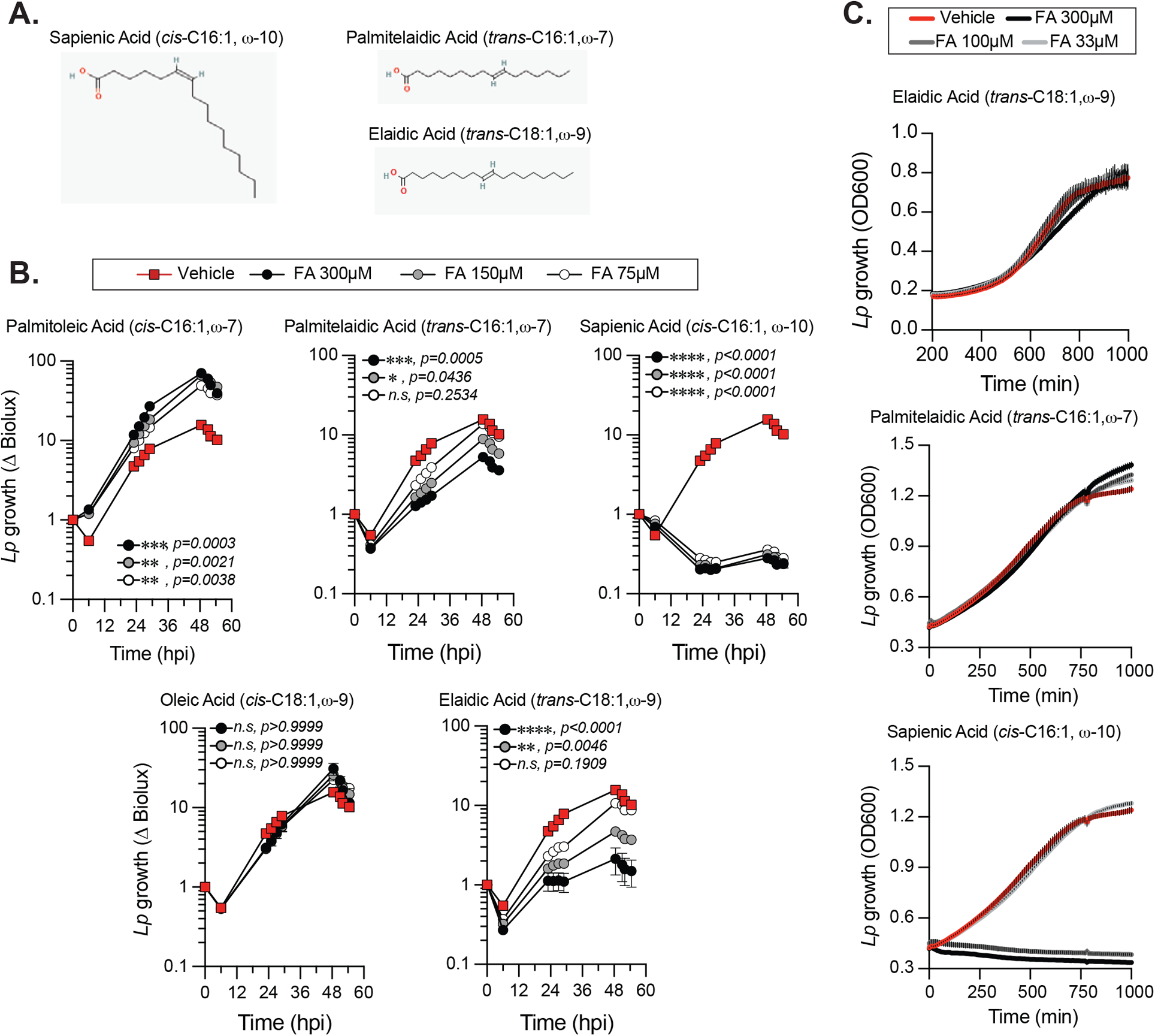
Effect of fatty acid positional and geometric isomerism on Lp growth in iBMMs. **(A)** Shown are the structures of (i) sapienic acid - a positional isomer of palmitoleic acid; (ii) palmitelaidic acid – a geometric isomer of palmitoleic acid and (iii) elaidic acid – a geometric isomer of oleic acid. Images were sourced from PubChem (pubchem.ncbi.nlm.nih.gov) **(B)** Lp growth kinetics with iBMMs at MOI=2.5 as measured by fold-change in bioluminescence signal from T_0_. The culture media was supplemented with the different fatty acids bound to BSA at the indicated concentrations or with BSA alone (vehicle control). Average ± StDev of three technical replicates are shown. Each data series from treated-cells was compared to vehicle-treated control using ordinary one-way ANOVA analysis and the respective p-values are indicated. At least three biological replicates for each experiment were generated. (*n.s* – not significant) **(C)** Axenic growth kinetics of Lp as measured by changes in optical density (OD_600_) over time in liquid cultures supplemented with increasing concentrations (33 to 300µM) of the indicated fatty acids bound by BSA or with BSA alone (vehicle control). OD_600_ data was collected every 10 minutes and each data series represents an average of three technical replicates ± StDev. **(B-C)** One representative of three biological replicates is shown for each experiment.

Robust microbiocidal activity towards Lp in axenic cultures (Fig 8C) and in macrophage infections (Fig 8B) was observed for sapienic acid - a non-linear positional isomer of palmitoleic acid in which the double bond is at the μ−10 carbon (Fig 8A). Sapienic acid is produced by Δ-6-desaturation of palmitic acid by FADS6 and is a major component of human sebum with antimicrobial properties (Ge et al., 2003; Drake et al., 2008). Thus, the position of the double bond on *cis*-C16:1 fatty acids can have remarkably different biological effects on *Legionella* viability and intracellular replication.

### Stearoyl-CoA desaturase – the enzyme desaturating palmitic (C16:0) to palmitoleic acid (C16:1) – is highly expressed by human pulmonary macrophages

The specificity of the Lp growth phenotype for palmitoleic acid over other fatty acids raised the possibility that palmitoleic acid might directly or indirectly interfere with cell intrinsic host defenses because palmitoleic acid can also function as a lipokine (Cao et al., 2008). As an adipose-tissue derived lipokine, palmitoleic acid regulates cellular responses through the activation of G protein-coupled receptors independently of membrane GPL remodeling. To investigate whether dietary palmitoleic acid broadly interferes with macrophage capacity to limit bacterial replication we decided to infect cells with other *Legionella* species that traffic and establish ER-like intracellular niches similar to *L. pneumophila* (MARUTA et al., 1998; Asare and Kwaik, 2007; Ivanov and Roy, 2009; Wood et al., 2015). To this end, we engineered *L. jordanis*, *L. longbeachae* and *L. dumoffii* strains that chromosomally encode the *luxR* operon under an IPTG-inducible promoter (SFig 5) and infected human U937 macrophages in the presence of palmitic or palmitoleic acid (Fig 9). Similar to the data from murine BMMs (Fig 1B-C and SFig 1A-B), palmitoleic acid but not palmitic acid significantly increased Lp growth in human macrophages demonstrating that the growth-promoting capacity of C16:1 is conserved (Fig 9). For the other *Legionella* species, palmitoleic acid did not enhance bacterial growth indicating that macrophages are not being affected in a manner that renders them broadly permissive for bacteria replication. Therefore, we conclude that the palmitoleic acid-induced growth advantage is conserved in human macrophages and is specific for *L. pneumophila*.

**Figure 9.**
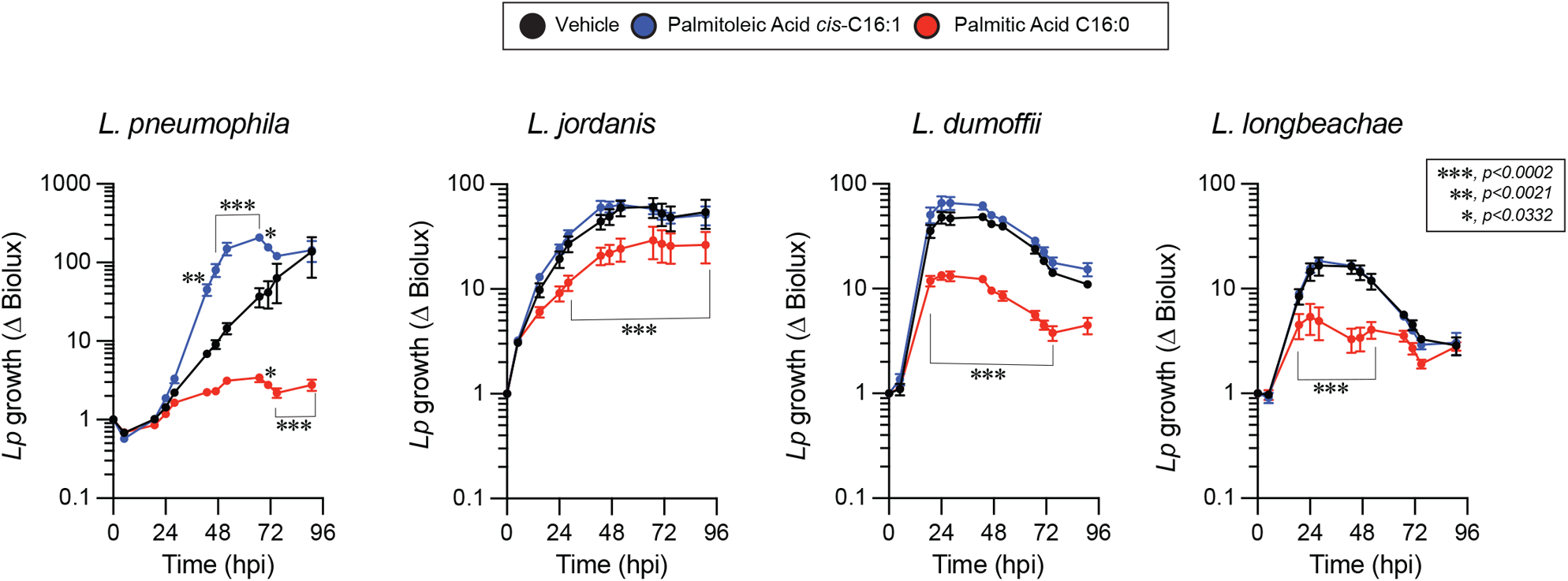
The growth-promoting effect of palmitoleic acid is specific for Lp in human macrophage infections. Growth kinetics of different *Legionella luxR* expressing species in human U937 macrophage infections in the presence of either BSA-complexed fatty acids (300µM) or BSA only vehicle control. Each data series represents an average of three technical replicates ± StDev. Each series was compared to vehicle control using two-way ANOVA analysis and time points with statistically significant p-values are indicated. Representative data from one of three biological replicates are shown.

Alveolar macrophages are a major cellular niche supporting Lp growth in human pulmonary infections (Nash et al., 1984), however they also perform a central role in the turnover of lung surfactant - a complex mixture of lipids, proteins and carbohydrates - that acts to decrease surface tension at the air–liquid interface of the alveoli in mammalian lungs (Bernhard, 2016). Type II alveolar epithelial (AE2) cells synthesize and secrete pulmonary surfactant, whereas both AE2 and alveolar macrophages carry out surfactant internalization and turnover (Stern et al., 1986; Lopez-Rodriguez et al., 2017). Phospholipids constitute ∼70% of pulmonary surfactant with dipalmitoyl-PC (PC16:0/16:0), palmitoyl-myristoyl-PC (PC16:0/14:0) and palmitoyl-palmitoleoyl-PC (PC16:0/16:1) being the major components (Bernhard, 2016). Thus, alveolar macrophages must be highly adapted to resolve the lipotoxic stress brought by continuous uptake and processing of GPLs containing palmitic acid. Eukaryotic cells cope with palmitate induced membrane rigidification by either (i) increasing the abundance of SCD – the enzyme that resolves membrane stress by desaturating palmitate to palmitoleic acid (Erbay et al., 2009; Piccolis et al., 2018; Oshima et al., 2020) or (ii) by transporting palmitate via the lipid chaperone Fatty Acid Binding Protein 4 (FABP4) to mitochondria for β-oxidation (Erbay et al., 2009). Gene expression profiles from single-cell RNAseq analysis of human lungs sourced from the Human Protein Atlas data repository (Karlsson et al., 2021) show very high levels of SCD expression in pulmonary macrophages and in AE2 cells, which would be needed to counteract membrane rigidification during surfactant turnover through SFA-to-MUFA conversion (Fig 10). The expression of *SCD5*, which encodes the other Δ9-desaturase enzyme allele in humans (Igal and Sinner, 2021) was low in both AE2 cells and macrophages (Fig 10). In addition, FABP4 transcripts were abundant in pulmonary macrophages but not AE2 cells consistent with the need for fatty acid transport to mitochondria for β-oxidation as part of surfactant turnover (Fig 10). Based on our data, palmitoleic acid sourced from surfactant-derived palmitoyl-palmitoleoyl-PC (PC16:0/16:1) is likely to increase the permissiveness of alveolar macrophages for Lp replication. Similarly, the high Δ9-desaturase activity in pulmonary macrophages would result in desaturation of surfactant-derived palmitic acid into palmitoleic acid and thus increase macrophage capacity to support Lp growth. Therefore, it is possible that the adaptive responses guarding against lipotoxic stress during the continuous turnover of pulmonary surfactant may inadvertently increase alveolar macrophages permissiveness for Lp replication.

**Figure 10.**
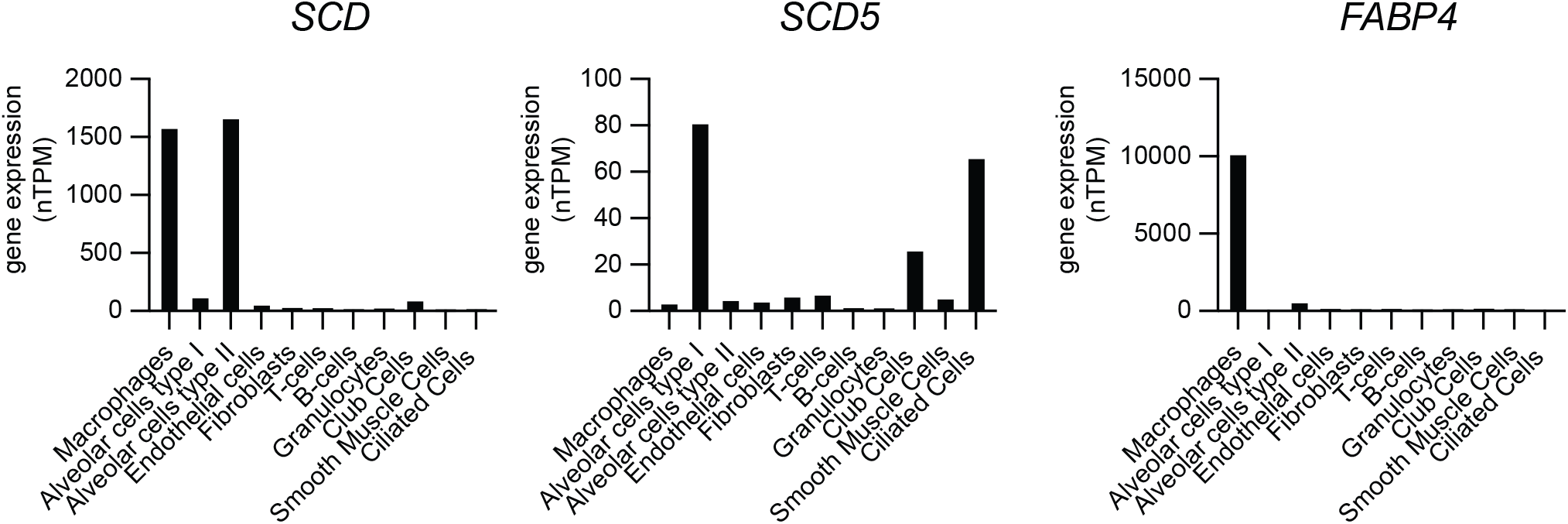
Gene expression analysis of *SCD*, *SCD5* and *FABP4* in different cell populations of the human lung. Single cell RNAseq gene expression data for *SCD* (Stearoyl-CoA desaturase), *SCD5* (Stearoyl-CoA desaturase 5) and *FABP4* (Fatty acid binding protein 4) in human lungs showing total transcript amounts as normalized protein-coding transcripts per million (nTPM) for each pulmonary cell type. Data is sourced from the Human Protein Atlas resource platform (proteinatlas.org) (Uhlén et al., 2015; Karlsson et al., 2021) – for SCD (www.proteinatlas.org/ENSG00000099194-SCD/single+cell+type/lung), SCD5 (www.proteinatlas.org/ENSG00000145284-SCD5/single+cell+type/lung) and FABP4 (www.proteinatlas.org/ENSG00000170323-FABP4/single+cell+type/lung).

## Discussion

This work provides experimental evidence that Lp replication within the LCV is a function of the acyl chain composition of the membrane glycerophospholipids. We took advantage of the preferential utilization of exogenous fatty acids by eukaryotic cells for membrane biogenesis to enrich specific acyl chains within the GPL repertoire and measured how this lipidome reprogramming impacted Lp intracellular replication. The data from a broad panel of saturated and monounsaturated fatty acids common in membrane GPLs demonstrated a surprising growth enhancement phenotype for palmitoleic acid in human and murine macrophage infections. The growth phenotype elicited by palmitoleic acid was associated with measurable effects on LCV homeostasis, such that (i) an increase in the number of LCVs that support bacterial replication; (ii) increased rate of bacterial replication; (iii) prolonged LCV expansion phase and (iii) shortened egress were observed. Combined, those changes significantly increased the number of bacteria produced from macrophage infections. Neither Lp uptake by macrophages nor Lp growth in axenic cultures was impacted by palmitoleic acid supplementation further supporting the notion that the growth-enhancing properties of palmitoleic acid are likely manifested through remodeling of host membranes. Indeed, our lipidomics data demonstrated that during infection upon palmitoleic acid treatment an extensive incorporation of C16:1 acyl chains was accompanied by a loss of C18:0 acyl chains across several GPL classes. Whether the ER membrane composition changes specifically under these conditions is a relevant question because Lp occupies an ER-derived organelle. The amount of multiple C16:1-acylated lipids from the two predominant ER-resident GPLs classes (PC and PE) increased dramatically upon palmitoleic acid treatment. Also, *de novo* GPLs synthesis (via the Kennedy cycle) begins with assembly of soluble precursors into phosphatidic acid by the sequential action of the ER resident transmembrane enzymes (glyceraldehyde-3-phosphate O-acyltransferase) (GPAT) and 1-acylglycerol-3-phosphate O-acyltransferase (AGPAT) (Jacquemyn et al., 2017). AGPATs also catalyze the re-acylation of lysophospholipids produced by PLA_2_ lipases in the Land’s cycle which results in direct replacement of acyl chain on preexisting GPLs (Shindou et al., 2013). Thus, regardless of whether C16:1 is incorporated in *de novo* produced or in preexisting GPLs the process is likely to take place at the ER because of enzyme localization.

Several lines of evidence point to membrane disorder as a critical LCV homeostasis variable. First, infections carried out in the presence of different MUFAs geometric isomers demonstrate that linear *trans*-isomers of C16:1 and C18:1 restricted Lp growth in macrophages without affecting the viability of the bacteria or the host cells, whereas the bend *cis*-isomers either enhanced (C16:1) or did not change (C18:1) Lp growth in macrophages. Second, *trans*-C18:1 treatment decreased the maximum LCV size as well as the rate of LCV expansion compared to the *cis*-C18:1 isomer consistent with the idea of membrane disorder benefiting LCV expansion. Both isomers were internalized comparably by macrophages and had similar effects on bacterial as well as host cell viability indicating that the phenotypic differences during infection are caused by other factors. Third, saturated fatty acids similar to *trans*-MUFA isomers restricted Lp growth albeit to various degrees. One-way LCV membrane disorder might benefit Lp is by affecting the function of the large repertoire of T4bSS transmembrane effectors (Dolezal et al., 2012) directly or by lowering the energy barrier for insertion of those effectors in the LCV membrane or by modulating membrane recruitment of Lp and host proteins (Vanni et al., 2014). Alternatively, increased access to small solutes across the LCV membrane might account for the enhanced Lp growth in an organelle with a high degree of membrane disorder because permeability in this case is a function of acyl chain saturation (Vanni et al., 2014). Membrane permeability might also explain the Lp growth enhancement caused by palmitoleic (C16:1) but not by oleic acid (C18:1) as increasing membrane hydrophobic thickness by elongating acyl chains by two carbons decreases permeability for passive diffusion of small solutes by a factor of ∼ 1.5 (Frallicciardi et al., 2022). Together, our data suggests that (1) niches with lipid phase ordered membranes generally support Lp replication poorly as compared to niches with disordered lipid bilayers and (2) decreasing hydrophobic thickness of the LCV might increase nutrient accessibility for small solutes passively diffusing across the LCV membrane.

The growth-enhancing capacity of palmitoleic acid for Lp in macrophage infections is particularly intriguing considering that alveolar macrophages continuously turnover pulmonary surfactant and thus function in a lipid-enriched environment. The high expression of the SCD desaturase by human pulmonary macrophages is likely a lipotoxicity-resolving adaptation that desaturates the C16:0 acyl tails, which predominate in surfactant lipids (Bernhard et al., 2001; Postle et al., 2006; Bernhard, 2016). Accumulation of C16:1 containing GPLs as a result of C16:0 desaturation within alveolar macrophages while resolving lipotoxic stress may inadvertently enhance macrophage capacity to support Lp replication as shown by our data and therefore would be detrimental during infection.

Multiple phospholipases produced by Lp and secreted through the T2SS (PlaA & PlaC) or the T4bSS (VipD and VpdC) have been shown to generate lysophospholipids through PLA_1_ or PLA_2_ enzymatic activity; therefore, those enzymes could potentially participate in remodeling of membrane GPLs. Due to their distinct subcellular localization different Lp PLA_1/2_ effectors might target separate lipid pools, where T4bSS effector lipases can access phospholipids in the cytosol-proximal leaflet and T2SS effectors hydrolyze lumen-proximal leaflet lipids. Regardless, it would be important to determine what substrates are hydrolyzed by Lp phospholipases when they are secreted intracellularly and whether these processes result in remodeling of the LCV membrane lipidome.

## Materials and Methods

### Bacterial Strains

All *L. pneumophila* strains used in this study were derived from the *L. pneumophila* serogroup 1, strain Lp01 (Berger and Isberg, 1993) and have a clean deletion of the *flaA* gene to avoid NLRC4-mediated pyroptotic cell death response triggered by flagellin when BMMs from C57BL/6J mice are infected with flagellated *Legionella*. The following Lp strains were used in this study: (1) *Lp01 ΔflaA*; (2) *Lp01 ΔflaA* Ptac-GFP expressing GFP under isopropyl-beta-D-thiogalactoside (IPTG)-inducible promoter; (3) *Lp01 ΔflaA LuxR* strain in which the *luxR* operon (*luxCDABE)* from *Photorhabdus luminescens* was inserted via homologous recombination on the bacterial chromosome downstream of the *icmR* promoter (Ondari et al., 2022). The following bioluminescent *Legionella* strains that encode the LuxR operon under the IPTG-inducible Ptac promoter on the chromosome were generated as described below: *L. dumoffii* strain NY 23, *L. jordanis* strain BL-540 and *L. longbeachae* strain Long Beach4. A list of the bacterial strains used in this study is provided in Table 2.

**Table 2.** Strains used in this study.

| Strains | Genotype | Properties | Reference |
| --- | --- | --- | --- |
| Lp01 $\Delta flaA$ | *L. pneumophila* serogroup 1, strain LP01 *rpsL*, $\Delta flaA$ | | (Ondari et al., 2022) |
| Lp01 $\Delta flaA$ *LuxR* | LP01 $\Delta flaA$, *icmR<sub>P</sub>-luxCDABE* | constitutive bioluminescence | (Ondari et al., 2022) |
| Lp01 $\Delta flaA$ GFP+ | LP01 $\Delta flaA$ Ptac-GFP | inducible GFP expression, *plasmid-borne* | (Ondari et al., 2022) |
| Lj *LuxR* | *Legionella jordanis* strain BL-540, Ptac-*luxCDABE* | inducible bioluminescence, *chromosome-borne* | This study |
| Ll *LuxR* | *Legionella longbeachae* strain Long Beach 4 (NCTC 11477), Ptac-*luxCDABE* | inducible bioluminescence, *chromosome-borne* | This study |
| Ld *LuxR* | *Legionella dumoffii* strain NY 23, Ptac-*luxCDABE* | inducible bioluminescence, *chromosome-borne* | This study |

*Legionella* strains were grown for 2 days at 37° C on charcoal yeast extract (CYE) plates [1% yeast extract, 1%*N*-(2-acetamido)-2-aminoethanesulphonic acid (ACES; pH6.9), 3.3mM l-cysteine, 0.33mM Fe(NO_3_)_3_, 1.5% bacto-agar, 0.2% activated charcoal] (Feeley *et al*., 1979). For all infections, each *Legionella* strain was harvested from CYE plates and grown in liquid cultures. For liquid cultures, bacteria from day 2 heavy patches grown on CYE plates were suspended to optical densities of 0.1 or 0.3 in 2.5ml of complete ACES buffered yeast extract (AYE) broth (10mg/ml ACES: pH6.9, 10mg/ml yeast extract, 400mg/l L-cysteine, 135mg/l) supplemented with 100µg/ml streptomycin and grown aerobically at 37°C to early stationary phase (between 18-22 hours, to OD 3.0-4.0). Liquid cultures of the GFP-expressing strains were supplemented with 10μg/ml chloramphenicol.

### Plasmids and strain construction

For the generation of inducible bioluminescent *Legionella* strains, genomic loci that contain an intergenic MfeI restriction site were chosen as follows: for *L. dumoffii* (Ld3X locus [315,471 _®_ 317,917 nt]); for *L. jordanis* (Lj5X locus [2,542,670 _®_ 2,545,129 nt]); for *L. longbeachae* (Ll7X locus [854,416 _®_ 856,719 nt]). The synthetic alleles that encode the *luxR* operon under the control of the tac promoter (Ptac) were built in stepwise fashion. **Step one** - the target genomic loci were PCR amplified from genomic DNA from each respective species with the following primers: Ld3X (*FC_BglII_MfeI003X* and *RC_SacI_MfeI003X*), Lj5X (*FC_BglII_MfeI005X* and *RC_SacI_MfeI005X*) and Ll7X (*FC_BglII_MfeI007X* and *RC_SacI_MfeI007X*). Each PCR amplicon product was double digested with BglII/SacI and cloned into the suicide plasmid pSR47s that was digested with BamHI/SacI. **Step two** - the Ptac regulon containing *lacI* (1,572 bp fragment) was PCR amplified using the pMR33 plasmid (Ramsey et al., 2012) as template with the primers *FC_EcoRI_Ptac regulon* and *RC_MfeI_Ptac regulon* and was cloned in the MfeI-digested pSR47s derivatives (from step 1) that contained the Ld3X, Lj5X and Ll7X loci. Correct directionality of the Ptac regulon within each locus was confirmed by sequencing. **Step three** - the pSR47s intermediates from step 2 were digested with BamHI and NotI and the *luxR* operon (5,854 bp fragment), which was similarly digested, was introduced to complete the construction of the pSR47s-Ld3X-Ptac-LuxR, pSR47s-Lj5X-Ptac-LuxR, and pSR47s-Ll7X-Ptac-LuxR plasmids.

The replacement of the target loci on the *Legionella* chromosome was carried out by allelic exchange via double homologous recombination and was confirmed via bioluminescence emission; kanamycin sensitivity and sucrose resistance (Merriam et al., 1997). For allelic exchange, plasmids were introduced in the respective *Legionella* species by either tri-parental mating (pSR47s-Ld3X-Ptac-LuxR and pSR47s-Lj5X-Ptac-LuxR) or via electroporation (pSR47s-Ll7X-Ptac-LuxR) with the following parameters voltage-1800, capacitance-25µF and resistance-200ohms. All primer sequences are listed in Table 3 and plasmid information is included in Table 4.

**Table 3.**
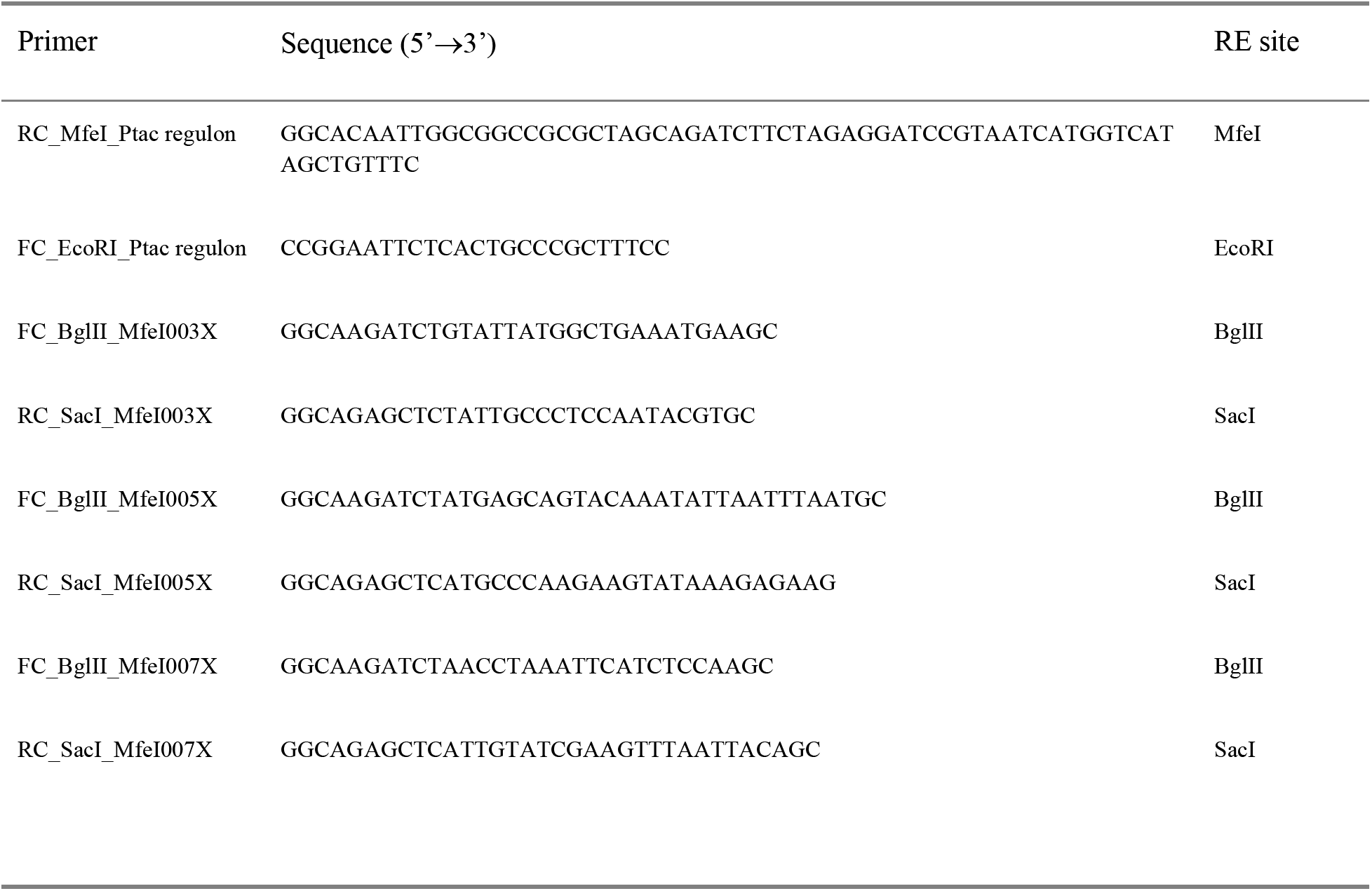
Primers used in this study.

**Table 4.** Plasmids used in this study.

| Plasmids | Important properties | Marker | Reference |
| --- | --- | --- | --- |
| pSR47s | Gene replacement vector | Kan | (Merriam et al., 1997) |
| pSR47s-Ld3X-Ptac-LuxR | pSR47s containing Ptac/luxR operon within the Ld3X locus (315,471→ 317,917nt) | Kan | This study |
| pSR47s-Lj5X-Ptac-LuxR | pSR47s containing Ptac/luxR operon within the Lj5X locus (2,542,670→ 2,545,129nt) | Kan | This study |
| pSR47s-Ll7X-Ptac-LuxR | pSR47s containing Ptac/luxR operon within the Ll7X locus (847,416→ 856,719nt) | Kan | This study |

### Reagents

The following reagents were purchased from Cayman Chemicals - lauric acid (cat# 10006626), myristic acid (cat# 13351), palmitic acid (cat# 10006627), palmitoleic acid (cat# 10009871), stearic acid (cat# 1011298), oleic acid (cat# 9004089), elaidic acid (cat# 90250), sapienic acid (cat #9001845), palmitelaidic acid (cat# 9001798), and Nile Red (cat#30787). Fatty acid-free Bovine Serum Albumin (BSA) was purchased from VWR (MP Biological, cat# 152401). Hoechst 33342 was purchased from ThermoScientific (Life Technologies, cat # H3570). The purified α-Lp IgY chicken antibody was custom generated by Cocalico Biologicals against formalin-killed bacteria (Abshire et al., 2016). The α-chicken-IgY-TRITC (cat#A16059) was purchased from ThermoFisher Scientific.

### Mice

C57BL/6J mice were purchased from Jackson Laboratories and housed at LSUHSC-Shreveport animal facility. Ethical approval for animal procedures for experiments in this study was granted by the Institutional Animal Care and Use Committee at LSUHSC-Shreveport (protocol# P-15-026).

### Macrophage cell culture conditions

Primary BMMs derived from bone marrow progenitors isolated from C57BL/6J mice were cultured on 10 cm petri dishes in RPMI 1640 with L-glutamine (BI Biologics, cat #01-100-1A) supplemented with 10% v/v FBS (Atlas Biologics, cat #FS-0500-AD), conditioned medium from L929 fibroblast cells (ATCC, CCL-1) (20% final volume) and penicillin/streptomycin (ThermoScientific, cat# 15140163) at 37°C with 5% CO_2_. Additional media was added at days 3, 5 and 7 from the start of differentiation. Macrophages were collected and seeded for infections at day 9. Immortalized BMMs from C57BL/6J mice were a kind gift from Dr. Jonathan Kagan (Harvard Medical School) (Evavold et al., 2021) and were propagated in High-Glucose DMEM supplemented with L-glutamine (ThermoScientific, cat# BP379-100), 10% v/v FBS (Atlas Biologics, cat# F-0500-DR), 1mM sodium pyruvate (Quality Biologicals, cat#116-079-721) and penicillin/streptomycin. U937 monocytes (ATCC, CRL1593.2) were cultured in RPMI-1640 with L-glutamine (BI Biologics, cat #01-100-1A), 10% v/v FBS and penicillin/streptomycin at a temperature of 37°C in the presence of 5% CO_2_.

### Complexing of different fatty acids with BSA

Fatty acid stocks were prepared in ethanol at the maximum possible concentration. Tissue-culture grade fatty acid-free BSA (MP Biomedicals, cat#152401) was dissolved in 150mM NaCl at a final concentration of 1.6mM and was filter sterilized through a 0.22µm PES filter. Fatty acid stock solution was mixed with the BSA stock solution in a 6:1 molar ratio to a final FA concentration of 10mM in 1.5ml Eppendorf tubes, which were subsequently incubated at 37℃ for 60min while being vigorously vortexed every 10min for at least 30sec. For the preparation of the BSA-only vehicle control stocks equivalent ethanol amount without the fatty acid was complexed with BSA.

### Cellular infections for lipidomics analysis

BMMs were seeded in 6 well tissue culture plates (2X10^6^ cells/well) in re-plating media (RPMI 1640 with L-glutamine, 10%FBS, 10% M-CSF-conditioned media) for 2hrs followed by 14hr incubation with serum-free RPMI (SF-RPMI). Cells were infected with liquid culture-grown *Lp01 ΔflaA* at MOI=5 for 8hrs in SF-RPMI. Plates were centrifuged for 5 min at 800rpms to increase bacteria attachment to host cells immediately after the inoculum was added. Palmitoleic acid complexed with BSA (200µM final concentration) was added at 2hpi in SF-RPMI. Each infection condition in every experiment was carried out in two technical replicates, which were pooled together after sample preparation. Samples from eight biological repeats were prepared for LCMS analysis as detailed below.

### Metabolite and Lipid Sample Preparation

For all LCMS methods LCMS grade solvents were used. Media was removed and discarded from cell culture samples in 6 well plates and washed briefly with 1 mL of 0.9 % sodium chloride. Wash was gently removed, and 0.4 mL of ice-cold methanol was added to each well. Plates were placed on ice for 5 min. To each sample 0.4 mL of water was added and the well was scraped to suspend cell-associated solids. Samples were transferred to microtubes and 0.4 mL of chloroform was added to each. Samples were agitated for 30 minutes at 4^°^C and subsequently centrifuged at 16k xg for 20 min to induce layering. An 0.4 mL aliquot of the organic layer (bottom) was dried in a Savant SpeedVac SPD130 (Thermo Scientific) and resuspended in 1 mL of 5 µg/mL butylated hydroxytoluene in 6:1 isopropanol:methanol.

### Liquid Chromatography Tandem Mass Spectrometry

LCMS grade water, methanol, isopropanol and acetic acid were purchased through Fisher Scientific. Bulk lipids were analyzed as previously described (Schwarz et al., 2020; Foliaki et al., 2023). A Shimadzu Nexera LC-20ADXR was used for chromatography across a Waters XBridge® Amide column (3.5 µm, 3 mm X 100 mm) with a 12-minute binary gradient from 100% 5mM ammonium acetate, 5% water in acetonitrile apparent pH 8.4 to 95% 5mM ammonium acetate, 50% water in acetonitrile apparent pH 8.0 was used to separate organic fraction samples. Lipids were detected using a Sciex 6500+ QTRAP® mass spectrometer with polarity flipping. Lipids were detected using scheduled MRMs based on acyl chain product ions in negative mode or a combination of invariant or neutral loss combinations in positive mode. All signals were integrated using MultiQuant® Software 3.0.3. Signals with greater than 50% missing values were discarded and remaining missing values were replaced with the lowest registered signal value. All signals with a QC coefficient of variance greater than 40% were discarded. Filtered datasets were total sum normalized prior to analysis. Single and multi-variate analysis was performed in MarkerView® Software 1.3.1. Datasets were z-scaled for display across the displayed groups. Z-score sums were calculated by summing all lipid signals with a given lipid characteristic in the z-scaled dataset that passed a statistical criterion. The entire lipidomics data set is available at Figshare via 10.6084/m9.figshare.24309139

### Microscopy analysis of lipids droplets formation in iBMM

Cells were seeded on coverslips in re-plating media (DMEM with L-glutamine, 10% FBS, sodium pyruvate) overnight at 3×10^5^ cells/well seeding density in 24-well plates. Cells were cultured in serum-free RPMI (SF-DMEM) for 6hrs and then were treated with either 48µM BSA alone or 150µM BSA-complexed fatty acid in SF-DMEM for 4hrs. Cells were gently washed with warm PBS (3X) and fixed with 2%-paraformaldehyde for 30min at ambient temperature. For quantitative analysis of lipid droplets, cells were stained with 200nM NileRed in PBS for 16hrs at 4°C and with Hoechst for 60min at ambient temperature. Coverslips were subsequently washed with PBS (5X) and mounted with ProLong Gold antifade reagent (ThermoFisher) onto glass slides. Images were captured with inverted wide-field Keyence BZX-800 microscope. Image acquisition parameters - Hoechst (ExW 395 / EmW 455) and NileRed (ExW 555 / EmW 605). The z-axis acquisition was set based on the out-of-focus boundaries and the distance between individual z-slices was kept at 0.3µm. Image analysis was performed with Keyence imaging software and the number of lipid droplets per cell was enumerated.

### Macrophage uptake assay

BMMs were seeded on coverslips as described in the previous section. Following attachment, cells were cultured in SF-RPMI containing either 300µM Palmitoleic acid complexed with BSA or BSA alone for 16hrs. Next, BMMs were infected with liquid culture grown *Lp01 ΔflaA* p*Tac*::GFP at MOI = 10 for 2hrs. One set of the BSA-treated cells was treated with 5µM cytochalasin D at 30min prior to the infection to block phagocytosis. After cells were washed with warm PBS (3X), the infection was stopped by the addition of 2% paraformaldehyde for 60min at ambient temperature. The surface-associated bacteria were immunolabeled without plasma membrane permeabilization with chicken α-*Legionella* IgY antibody for 90 min in PBS containing goat serum (2% vol/vol). Next, coverslips were washed with PBS (3X) and were stained with tetramethyl rhodamine conjugated goat α-chicken IgY (ThermoFisher, cat# A16059) at 1:500 dilution and Hoechst at 1:2000 dilution for 60min in PBS containing goat serum (2% vol/vol). Coverslips were mounted with ProLong glass anti-fade mountant (ThermoFisher, cat# P36984) onto slides.

Microscopy analyses of infected cells. Images were captured with inverted wide-field Keyence BZX-800 microscope. Image acquisition parameters - Hoechst (ExW 395 / EmW 455); GFP (ExW 470 / EmW 525) and TRITC (ExW 555 / EmW 605). The z-axis acquisition was set based on the out-of-focus boundaries and the distance between individual Z-slices was kept at 0.3µm. Only linear image corrections in brightness or contrast were completed. For each condition, over 100 bacteria were imaged and scored as either intracellular (single positive – green only) or not-internalized (double positive – green/red).

### *Legionella* axenic growth assays

Liquid cultures of *Lp01 ΔflaA LuxR* in complete AYE were set-up at starting OD_600_ = 0.4 from plate grown bacteria (day-2 heavy patches) and were distributed in white-wall clear-bottom 96-well plates (Corning, cat# 3610). BSA-complexed fatty acids were added at the beginning of the assay at the indicated concentrations. All conditions in all assays were performed in technical triplicates. Plate was incubated in a luminometer (Tecan Spark) at 37°C for 24 hrs. Optical density (OD = 600 nm) data were automatically collected every 10 mins after the cultures were agitated for 180sec (double orbital rotation, 108 rpms).

### Bioluminescence assay for *Legionella* intracellular replication

Primary BMMs were seeded at 1×10^5^ cells per well in white-wall, clear bottom 96-well plates (Corning cat# 3610) in re-plating media (RPMI 1640 with L-glutamine, 10% FBS, and 10% M-CSF-conditioned media) for 2hrs. Immortalized BMMs were seeded in re-plating media of High Glucose-DMEM with L-glutamine (5.5mL of 100X, ThermoScientific Cat# BP379-100, FBS (10% final volume, Atlas Biologics, cat# F-0500-DR), sodium pyruvate (5.5mL of 100mM, Quality Biologicals Cat# 116-079-721, and penicillin/streptomycin (5.5mL of 100X, ThermoScientific Cat# 15140163). For human U937 macrophages infections, U937 monocytes were seeded at a density of 1 × 10^5^ cells per well in white-wall clear-bottom 96-well plates and differentiated into macrophages after 24hr treatment with 10ng/ml Phorbol 12-myristate 13-acetate (Adipogen) after which cells were cultured without PMA and antibiotics for additional 48hrs prior to infection. Cells were cultured with serum-free media for 14hrs (BMMs and U937 macrophages) or for 6hrs (iBMMs) prior to infection. Infections were carried out under serum free conditions with liquid culture grown *Lp01* Δ*flaA icmRp-LuxR* at MOI=5. BSA-complexed fatty acids were added at 2hpi at the indicated final concentration. Plates were kept in a tissue culture incubator at 37°C with 5% CO_2_ and periodically, the bioluminescence output from each well was acquired (integration time 2 or 5sec). The data is presented as fold change in bioluminescence from the T_0_ reading. All conditions in all assays were performed in technical triplicates.

### Colony Forming Units assay for *Legionella* intracellular replication

Macrophages were seeded, infected, and treated as listed in “*Bioluminescence assay for Legionella intracellular replication*”. Plates were kept in a tissue culture incubator at 37°C and 5% CO_2_ and periodically removed to collect bacteria. For bacterial recovery, cell culture media from the well was moved into a 1.5ml Eppendorf tube and replaced with 100µL of sterile water for lysis of the infected macrophages via hypotonic shock and the plate was returned to the incubator for 10mins. Next, the contents of the well were pipetted at least 20 times and were collected in the respective Eppendorf tube which already contained the cell culture media. Eppendorf tubes were vortexed for 30sec and contents (∼300µL) were serially diluted five times for a total of six dilutions. 25µL from each dilution was plated on a CYE agar plate and incubated until colonies were visible. Colonies from a single dilution were counted and CFUs were calculated for each condition. All infection conditions for every experiment were performed in three technical replicates. The data is presented as fold change in recovered CFU over T_0_.

### Microscopy analysis of *Legionella* intracellular growth and calculation of the Replication Vacuole Index

iBMMs were seeded on coverslips at 3×10^5^ cells per coverslip in re-plating media (DMEM with L-glutamine, 10% FBS, sodium pyruvate). After overnight incubation at 37°C and 5% CO_2_, cells were cultured with serum-free media and infected with *Lp01 ΔflaA* GFP+. At 2hpi, infections were gently washed with warm PBS (3X) to remove extracellular bacteria. One infection condition was stopped at 2hpi, the remaining infections were allowed to proceed until 10hpi in the presence of serum-free media containing either BSA only, 300µM palmitoleic acid or 300µM palmitic acid. At 10hpi, cells were washed with warm PBS (3X) and the infection was stopped by the addition of 2% paraformaldehyde for 60min at ambient temperature. Next, coverslips were stained with Hoechst (1:2000 dilution) for 60min and were mounted with ProLong glass antifade mountant (ThermoFisher, cat# P36984) onto slides. The number of infected cells and the number of bacteria per LCV were enumerated for every condition. The Replicative Vacuole (RV) Index is calculated as the ratio of the percentage of infected macrophages at 10hpi that harbor LCV with >6 bacteria over the percentage of infected macrophages at 2hpi.

### IncuCyte^TM^ S3 automated microscopy analysis of *Legionella* intracellular replication

For infections, BMMs were seeded in 96-well black-wall clear-bottom plates (Corning cat# 3904) at 8.0×10^4^ cells per well in re-plating media (RPMI-1640 with L-glutamine, 10% FBS, and 10% M-CSF-conditioned media) for 2 hours and then were serum starved for 14hrs in serum-free phenol red-free DMEM (Gen Clone 25-501C). All infections were carried out in serum-free media supplemented with 1mM IPTG. BSA-complexed fatty acids were added at 2hpi. Final media volume was 150µl per well. In each experiment, all conditions were performed in technical triplicates. Plates were centrifuged (1,000rpms, 5min) and were loaded into the IncuCyte^TM^ S3 housing module. The IncuCyte^TM^ S3 HD live-cell imaging platform (Sartorius) is a wide-filed microscope mounted inside a tissue culture incubator and is run by the IncuCyte control software. For each well, four single plane images in bright field and green (ExW 440-480nm/EmW 504-544nm) channels were automatically acquired with *S* Plan Fluor 20X/0.45 objective every four hours. Images were analyzed with the IncuCyte Analysis software. For LCV analysis, bacterial fluorescence was used to generate a binary mask to define individual LCV objects and to measure the object’s area size (GCUxµm^2^) and the number of objects per image. For single object LCV tracking, the LCV area was recorded for ∼100 LCVs per condition for every timeframe from appearance of the LCV to its disappearance. The IncuCyte^TM^ S3 imaging analysis was performed in the Innovative North Louisiana Experimental Therapeutics program (INLET) core facility at LSU Health-Shreveport.

### Measurement of cell viability

BMMs were seeded at 1×10^5^ cells per well in white-walled, clear bottom 96-well plates (Corning cat# 3610) in re-plating media (RPMI 1640 with L-glutamine, 10% FBS, and 10% M-CSF-conditioned media) until attached. Macrophages were serum-starved overnight prior to treatment with different fatty acids. Cell viability was assessed continuously based on ATP production with the RealTime-Glo^TM^ MT Cell Viability Assay kit (Promega, cat# G9711) through bioluminescence output using the TECAN Spark plate reader.

### Analysis of pulmonary gene expression

Single cell RNAseq data from normal human lung samples for SCD, SCD5 and FABP4 was sourced from The Human Protein Atlas resource platform (https://www.proteinatlas.org/)(Uhlén et al., 2015; Karlsson et al., 2021). For each transcript the original sourced data is presented in normalized protein-coding transcripts per million (nTPM). For each cell group the transcript expression data is pooled from all representative cell types. The original gene expression data can be found through these links: SCD (www.proteinatlas.org/ENSG00000099194-SCD/single+cell+type/lung); SCD5 (www.proteinatlas.org/ENSG00000145284-SCD5/single+cell+type/lung); FABP4 (www.proteinatlas.org/ENSG00000170323-FABP4/single+cell+type/lung).

### Statistical analysis

Calculations for statistical differences were completed with Prism v10.0.3 (GraphPad Software). The statistical tests applied for the different data sets are indicated in the figure legends and the resultant p-values are shown in the figures.

---

**Supplemental Figure 1. Exogenous fatty acids alter Lp growth kinetics in iBMMs infections. (A-B)** Lp growth kinetics with iBMMs at MOI=2.5 as measured by fold-change in bioluminescence signal **(A)** or recovery of colony forming units **(B)** from T_0_. The culture media was supplemented with different fatty acids bound to BSA at the indicated concentrations or with BSA alone (vehicle control). **(A)** Average ± StDev of three technical replicates are shown. Each data series from treated-cells was compared to vehicle-treated control using ordinary one-way ANOVA analysis and the respective p-values are indicated. At least three biological replicates for each experiment were generated. **(B)** Graph shows CFUs data recovered at 48hpi from four biological replicates, which were analyzed for statistical significance with a ratio-paired *t*-Test.

**Supplemental Figure 2. Live-cell imaging of LCV expansion.** From live-cell imaging of primary BMMs infections with GFP-expressing Lp, the graphs show individual tracers tracking the change in size over time for ten randomly selected LCV for each of the indicated conditions from the data presented in Fig 8. The size is measured by the total integrated green fluorescent intensity (GCU x μm^2^) for each object.

**Supplemental Figure 3. Time-lapse microscopy analysis of LCV collapse.** Live-cell imaging of Lp intracellular replication in primary BMMs infected with GFP-expressing bacteria at MOI=5. BSA-complexed exogenous fatty acids (150µM) were added at 2hpi. The length of LCV collapse phase – from the end of the stasis phase until complete disappearance - (as defined in Fig 8a) is graphed for each condition based on the analysis of multiple individual LCVs. The number of LCVs analyzed for each condition is indicated.

**Supplemental Figure 4. Cell viability in fatty acid-treated pBMMs. (A-B)** Changes in cell viability over time of primary BMMs treated with either the indicated BSA-complexed fatty acids (300µM) or BSA only (vehicle) were measured continuously over the indicated time period with Promega’s RealTime-Glo^TM^ MT Cell Viability assay. Treatments with saturated and monounsaturated fatty acids are shown in **(A)** and **(B),** respectively. Each data series represents an average of three technical replicates ± StDev. For statistical analysis, each data series was compared to the data series obtained from the vehicle-treated cells using a one-way ANOVA analysis and the respective p-values are indicated. Representative data from one of two biological replicates are shown.

**Supplemental Figure 5. Genomic organization of the *luxR* locus in the different *Legionella* species used in this study.** The *LuxR* operon from *Photorhabdus luminescens* under the control of the P_TAC_ promoter was introduced in an intergenic region on the chromosome of the indicated *Legionella* species via allelic exchange as is shown in each schematic.

## Acknowledgments

We would like to thank Prof. Jonathan Kagan (Harvard Medical School) for providing the immortalized C57BL/6 bone marrow-derived macrophage cell line and the INLET High-Throughput Imaging Core at LSUHSC-Shreveport for technical assistance.

## Data Availability Statement

The lipidomics datasets generated for this study can be found in the Figshare via 10.6084/m9.figshare.24309139.

