## Supplementary figures and images for "The intracellular growth of the vacuolar pathogen *Legionella pneumophila* is dependent on the acyl chain composition of host membranes"

### Supplemental Fig 1

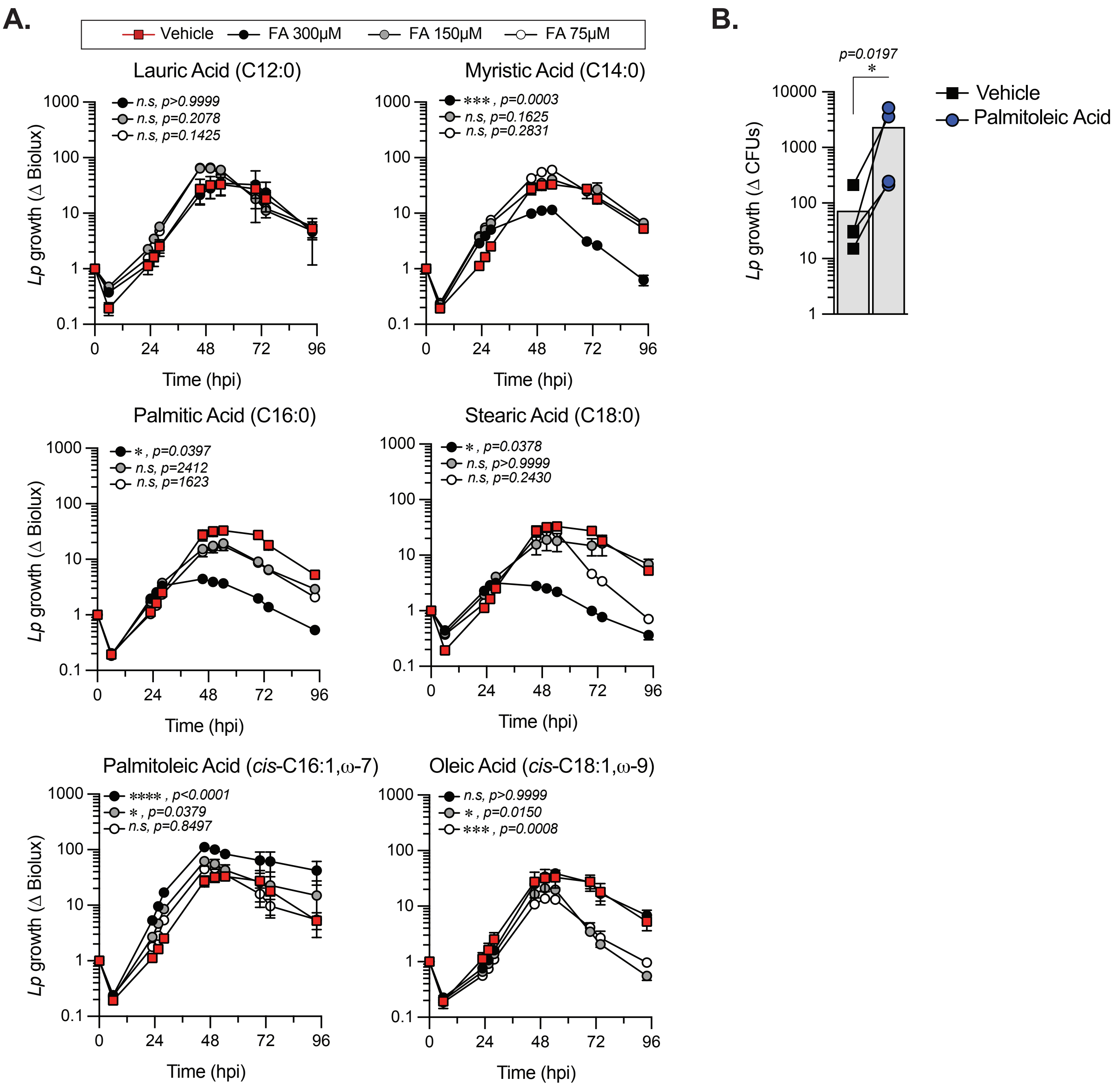

### Supplemental Fig 2

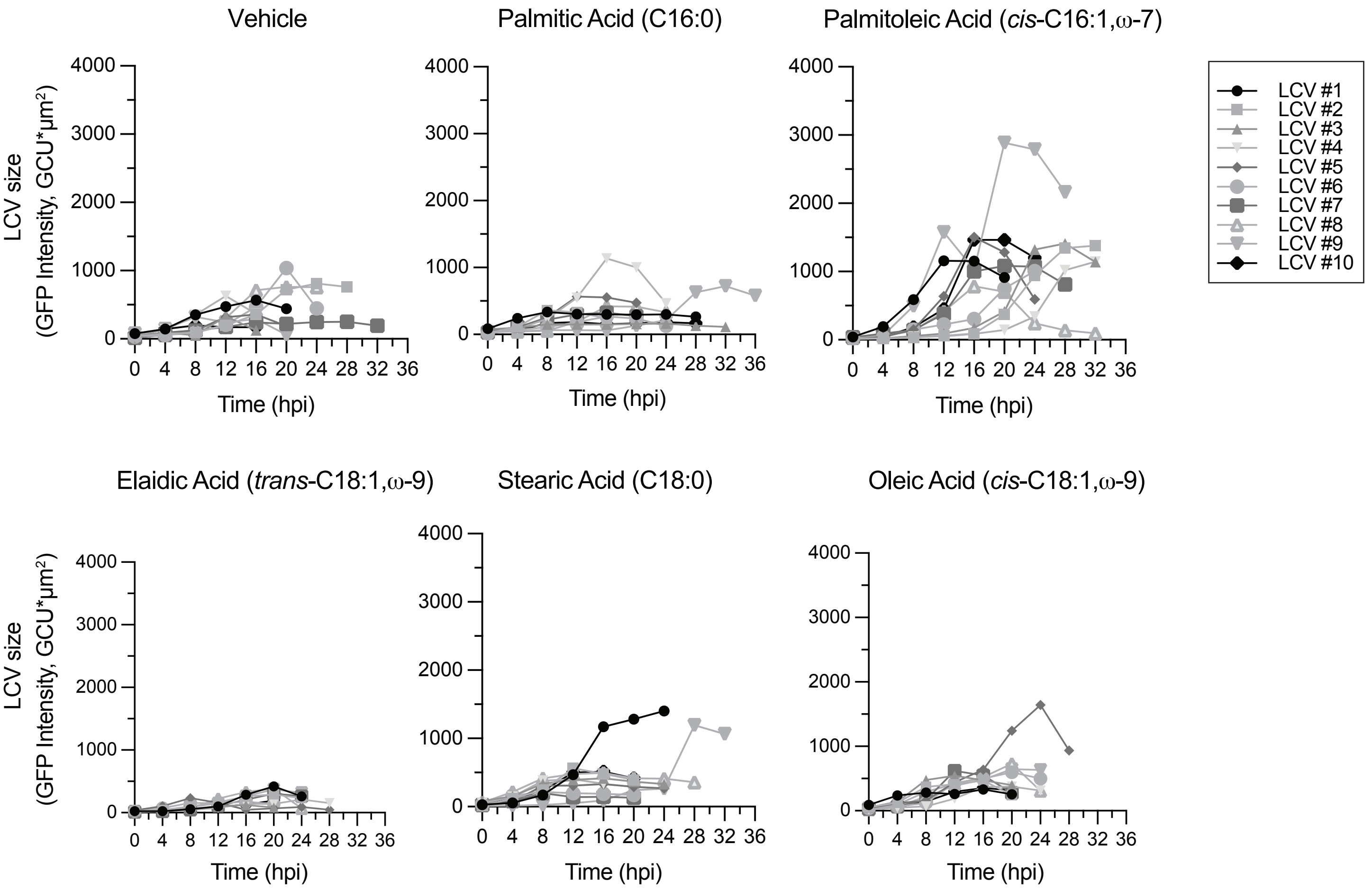

### Supplemental Fig 3

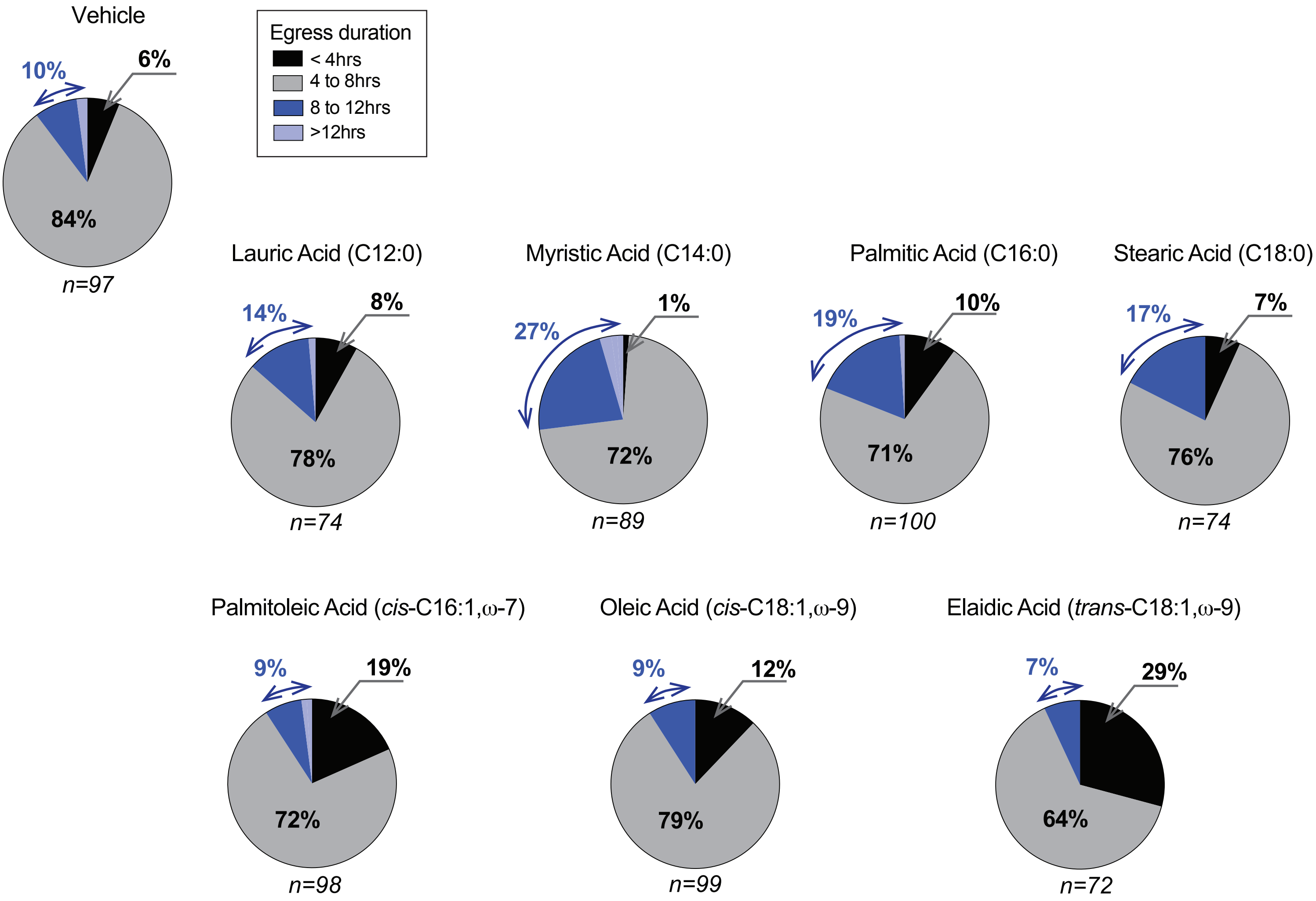

### Supplemental Fig 4

#### Sup. Figure 4.

**A.**

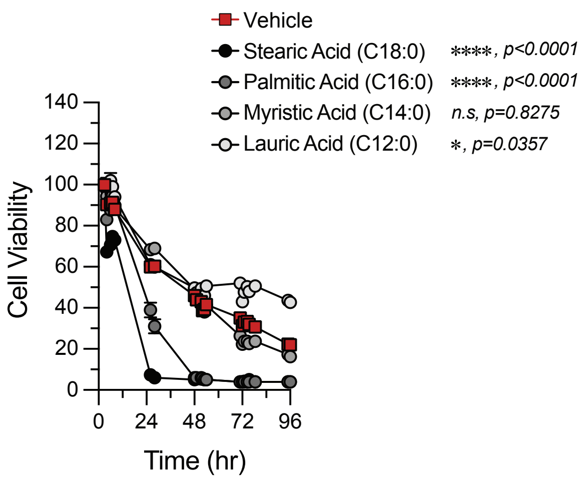


**B.**

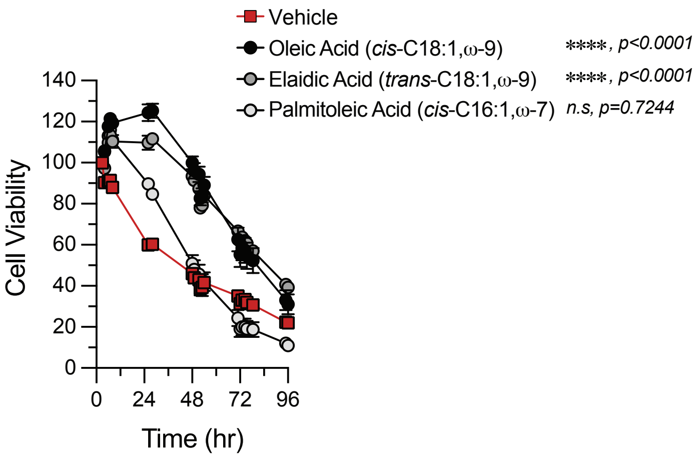

### Supplemental Fig 5

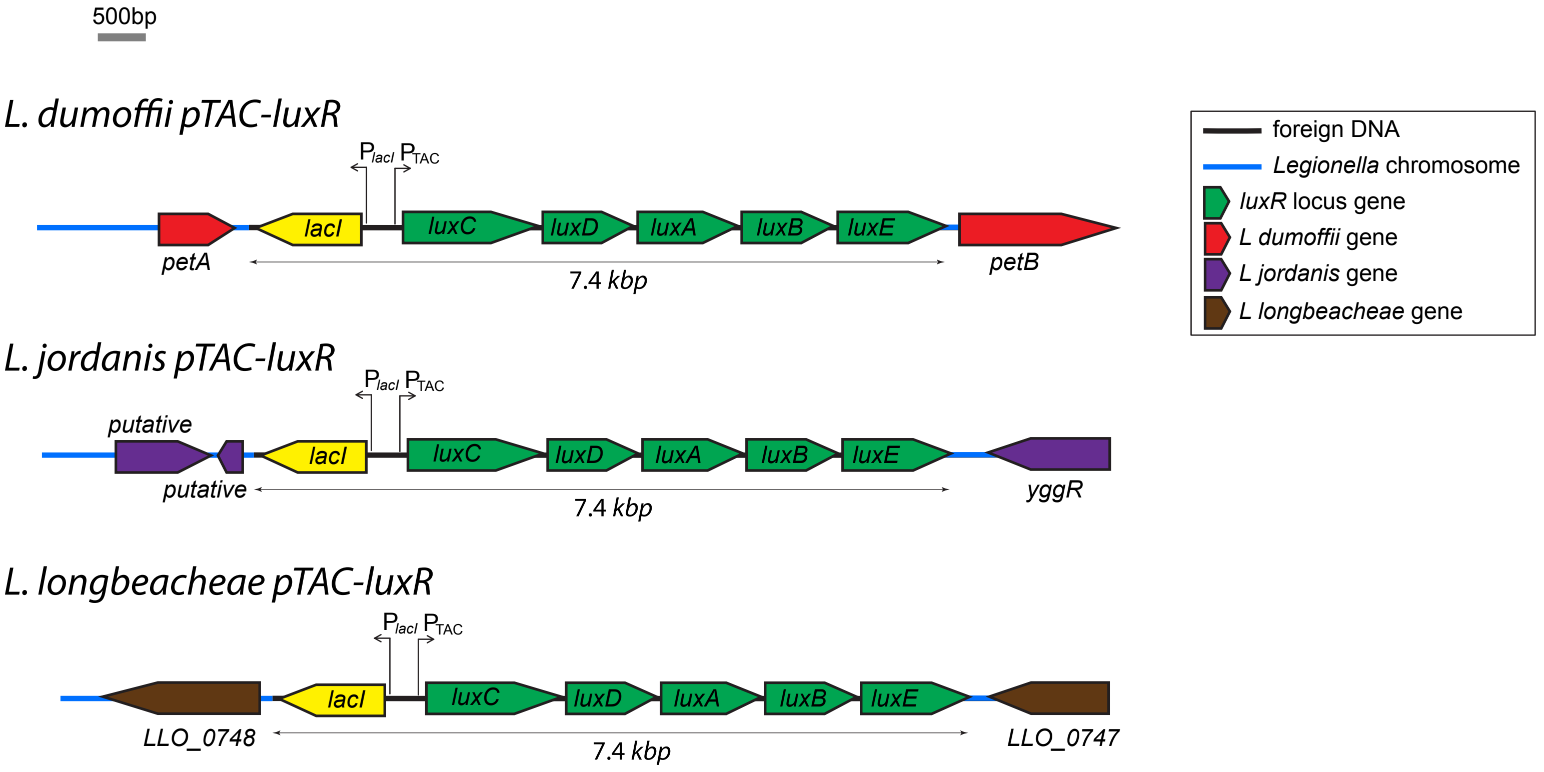
